# Clade-specific influences of glycans on the interactions between HIV-1 envelope and broadly neutralizing antibodies

**DOI:** 10.1101/2025.05.20.655089

**Authors:** Mrinal Arandhara, Yogendra Kumar, Narendra M. Dixit, Prabal K. Maiti

## Abstract

N-linked glycans are important in the elicitation and the activity of many broadly neutralizing antibodies (bNAbs) against HIV-1. The high conformational flexibility of glycans hindered detailed atomistic investigations of glycan-bNAb interactions, including glycan shielding of bNAbs. Importantly, how these interactions vary across different HIV-1 clades remains unclear. The variability in the number and location of potential N-linked glycosylation sites (PNGS) on the HIV-1 envelope (Env) protein across clades can lead to differences in glycan dynamics and topology, potentially affecting Env-bNAb interactions and the clade-specific efficacy of bNAb-based therapies. Here, we combined comprehensive glycan conformational sampling, using the software glycoSHIELD, and molecular dynamics (MD) simulations to model fully glycosylated trimeric Env for six HIV-1 strains, one from each of the major clades A, B, C, G, CRF01 AE (01 AE) and CRF07 BC (07 BC). We assessed the interactions of 50 different bNAbs, drawn from all the major bNAb classes, with each of these strains, quantifying glycan shielding, glycan-bNAb interactions, and their clade-specific variations for each bNAb in microscopic detail. Our findings reveal that while glycans cover most of the exposed surface area in all clades, the amount of accessible surface varies, with clade B having the minimum and clade 07 BC the maximum antibody accessible surface area. The number of glycan conformers per glycosylation site also varies with clades, even for conserved sites. Overall, we observed that bNAbs interact with more glycans than previously reported in experimental and computational studies. Important variations emerge in Env-bNAb interactions with clade and bNAb-class. These atomic-level insights will be valuable for improving bNAb-based therapies and vaccine design strategies against HIV-1.

**TOC Graphic:** 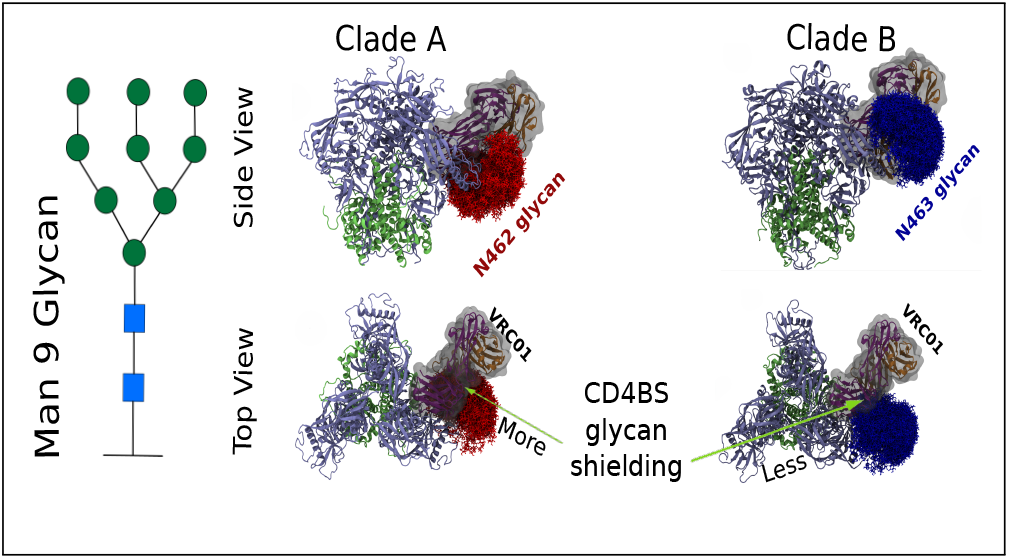

## Introduction

The envelope spike (Env) protein on the HIV-1 virion surface is the only target for neutralizing host immune responses.^1,2^ The viral spike consists of a trimer of gp120 and gp41 heterodimers, where gp120 and gp41 in each monomer are held together through non-covalent interactions.^3^ Compared to other glycoprotein viruses such as SARS-CoV-2 and influenza, the HIV-1 Env is heavily glycosylated.^4–7^ Nearly half of the total mass of the Env is due to surface glycans.^8–10^ These glycans protect the virus from host immune responses,^11^ contribute to Env’s structural stability,^12,13^ and maintain the native folding of the Env.^14^ As native-like structure is vital for eliciting neutralizing antibodies, glycans are essential for vaccine design.^15^ Despite the challenges of overcoming the glycan barrier, a fraction (≈1 ™ 2 %) of people living with HIV-1, called elite neutralizers, develop highly potent antibodies, called broadly neutralizing antibodies (bNAbs), which can effectively neutralize diverse strains of HIV-1. ^16–18^ Most bNAbs have unique features, such as high somatic hypermutation and an unusually long CDR3 loop on the heavy chain, that allow them to navigate the glycan shield and bind to the viral protein^1,19^. Moreover, all the known bNAbs are found to engage with at least one glycan on the Env^10^. Except for the bNAb 2G12, which only targets glycans on the viral surface, all bNAbs target the Env surface along with the glycans. ^11^

HIV-1 is typically divided into three groups: Group M (major), N (non-major), and O (outlier). Group M is the most prevalent, infecting more than 90% of people living with HIV^20,21^. Group M is further divided into subtypes or clades, with different clades prevalent in different geographical locations. Clade A is predominant in West and Central Africa^21,22^, clade B in America and Western Europe ^21^, and clade C in Southern Africa, India, and parts of China^21^. In addition, circulating recombinant forms, such as 01 AE and 07 BC, dominate in Southeast and East Asia^21,23^. Vaccines that can target these diverse sets of HIV-1 viruses are much needed.

Based on the specific regions on the Env they target, bNAbs are broadly classified into the following categories ^1,16,19^: CD4 binding site (CD4bs), V1/V2 apex, V3-glycan, gp120-gp41 interface, fusion peptide (FP), and membrane-proximal external region (MPER) (see Figure S1 in the SI). bNAbs targeting the V1/V2 apex (e.g., PG9, PG16, PGT145) and V3-

glycan (e.g., PGT121, 10-1074) require glycans for recognition or neutralization. Glycans form a key epitope of many V1/V2 apex and V3-glycan targeting bNAbs^1,11,15,19^. Although CD4bs-targeting bNAbs generally do not require glycans for recognition, their structural features allow them to accommodate or evade glycans at or near the epitope sites. ^24,25^

Numerous studies have investigated the potential use of bNAbs and dendrimers for prevention, treatment, and cure of HIV-1. ^1,26–31^ A mechanistic model was developed to elucidate how passive immunization (PI) with high-affinity antibodies can potentiate endogenous antibody production. The model integrates diverse experimental observations and offers strategic insights for optimizing PI protocols to enhance immunological outcomes.^32^ An early passive immunization with bNAbs can lead to long-term viral control in SHIV-infected macaques. Mathematical model of viral dynamics demonstrates that early bNAb therapy can shift the infection outcome from progressive disease characterized by high viremia and immune exhaustion to a state of lasting control with low viremia and robust immune responses.^33^ HIV-1 variants are resistant to VRC01 can exist in the latent reservoir—a pool of dormant infected cells unaffected by ntiretro-viral therapy (ART). Upon cessation of ART and administration of VRC01, these resistant strains can become reactivated and undergo replication leading to therapeutic failure. ^34,35^

In pre-fusion of HIV-1, the gp41 interacts with different lipids in the membrane and forming stalk while in simpler lipid membrane shows strong resistance to fusion. ^36^ However, in the post-fusion process, the gp41 interacts strongly with C34 and minimally with T20 peptide inhibitors.^37^ Many clinical trials involving bNAbs have been completed and many are cur-rently ongoing. Efforts are also underway to develop vaccines that elicit these antibodies^1,11,19,26,27,38,39^ The dynamics of affinity maturation within germinal centers was strategically modulated to elicit bNAbs against HIV. ^40^ In contrast, a sequential immunization strategy, presenting antigen variants one at a time, promotes the targeting of conserved regions such as the CD4 binding site.^40^ Although they can be as effective as antiretroviral treatments in maintaining viral loads, bNAbs often have better half-lives and/or toxicity profiles, making them promising alternatives for future treatment^27^. They also seem to lead to better long-term remission post-treatment than antiretroviral therapies, the reasons for which are beginning to be elucidated. ^41,42^ However, efforts to induce bNAbs through vaccination have largely been unsuccessful^11,19^.

Recent studies have made significant progress in characterizing the glycan shield and understanding glycan-bNAb interactions for a few bNAbs. Env glycans are mostly N-linked and occur at the consensus sequence Asn-X-Ser/Thr, where X is any amino acid (AA) other than proline^43^. Site-specific mass spectrometry studies revealed that Env glycans are composed of three types of glycans, termed high-mannose, complex, and hybrid^4,5,44–47^. Table S1 and Figure S2 in the SI show schematics and 3D structures of various glycans, respectively. The glycan type is site-dependent and typically determined by glycan crowding around it, with more crowded glycans being less processed^6,10^. Numerous experimental and theoretical studies have also yielded insights into glycan-bNAb interactions^1,11,15,19,24,25,38,48–53^. Molecular dynamics (MD) simulations of glycosylated clade A strain has been key in understanding glycan dynamics and their influence on a few bNAbs for^24,25,54–56^. Yet, the high degree of conformational flexibility and complexity of glycans has hampered both experimental and computational investigations^53,57^. Due to the highly dynamic nature of the glycans, their structures are not correctly resolved in the cryoelectron microscopy experiments^14,53,57^. The long time scale required to sample the relevant glycan conformations and the numerous glycosylation sites make MD simulations challenging^53,55,57^. Using cryo-EM data and a computational approach, Berendsen *et al*.^14^ quantified and visualized glycan shielding but only for clade A structure BG505 SOSIP.664. Wagh *et al*.^58^ utilized a sequence-based approach to approximately model glycan shielding of an ensemble of HIV-1 strains, but their approach lacks atomistic details.

Most HIV-1 research has focused on a few clades, specifically B and C, with limited knowledge, therefore, of other clades available. Clade-specific differences can be significant.^59^ Rusert *et. al*. observed that the types of bNAb elicited in people with HIV-1 are influenced by the HIV-1 clade^60^. The number of Env glycans can vary across clades , leading to variation in glycosylation patterns on the surface. Besides, clade-specific variation in the local protein structure surrounding a glycan site can alter the structure and dynamics of glycans. In addition to other parameters, such as AA sequences, these factors may influence the effectiveness and sensitivity of bNAbs.

In this work, using the software GlycoSHIELD (GS)^57^ in combination with MD simulations, we have generated fully glycosylated trimeric Env and investigated the variation in glycan shields and their impact on Env-bNAb interactions for 50 bNAbs and all the major HIV-1 clades. Specifically, we chose one strain from each of the clades A (strain:BG505), B (JR-FL), C (strain: CH848.10.17), G (strain: X1193.c1), CRF01 AE (strain: BJOX018000.e12), and CRF07 BC (strain: BJOX016000.e15). GS allows the generation of realistic models of glycoproteins and has been shown to reproduce experimental and MD simulation results on protein shielding and epitope accessibility mediated by glycans accurately^57^.

The manuscript is organized as follows: First, we describe the computational methodologies used. Next, we present the results of our analysis focusing on how glycosylation patterns vary across clades and then on how it affects binding to various bNAbs, highlighting the main findings and their implications. Finally, we conclude with a summary of key observations and future outlook.

## Methods

### Modelling of trimeric HIV-1 Env

The analysis performed in this work requires 3D trimeric structures of the HIV-1 Env protein. As a first step toward building these models, we collected high resolution structures (x-ray difraction or cryo-EM) of the Env trimer from the Protein Data Bank (www.RCSB.org) .^61^ Specifically, we took one representative crystal structure for each of the following six clades: A (strain: BG505, PDBID 5FYL),^24^ B (strain: JR-FL, PDBID 5FYK),^24^ C (strain:CH848, PDBID 8SAU), ^62^ G (strain: X1193.c1, PDBID 5FYJ),^24^ and circulating recombinant forms 01 AE (strain: BJOX018000.e12, PDBID 8GPI)^23^ and 07 BC (strain: BJOX016000.e15, PDBID 8GPJ)^23^. Missing amino acid residues were modeled using the DOPELoopModel method in the Modeller10.5 software.^63^ We generated five structural models for each clade and selected the structure with the best (lowest) DOPE score. We added H-atoms to the selected structures using Gromacs (version 2019.6)^64^ software. Next, we took 11 additional geometries from a 10 ns long MD simulation of each of the selected structures as discussed below. Figure1 displays the modeled structures for clades A, B, C, G, 01 AE, and 07 BC.

### Modelling bNAbs

We selected crystal structures of 50 bNAbs belonging to six different classes: CD4bs, V1/V2 apex, V3-glycan, gp120-gp41 Interface, fusion peptide, and glycan. We chose only those bNAbs bound to a domain of HIV-1 Env protein in the crystal structure. All structures were obtained from the rcsb database (www.RCSB.org).^61^ bNAbs targetting the MPER and silent face are not considered here due to the unavailability of reliable HIV-1 bound crystal structures. Figure S1 shows the trimeric Env structure of clade B (strain: JR-FL, PDBID: 5FYK) [left: Top View, Right: Side View] with the locations of various epitope classes: CD4bs Ab epitope (red), V1/V2 Ab epitope (green), V3 loop Ab epitope (blue), fusion peptide (FP) Ab epitopes (orange), and gp120-gp41 interface Ab epitope (Purple). Table S2 contains the list of bNAbs with associated PDB codes. bNAb structures with only the Fv portion are shown in blue in Table S2.

### Generation of glycoprotein trajectories using GlycoSHIELD

The molecular complexity and flexibility of glycan and the presence of many PNGS on HIV-1 trimer make long-time MD simulation challenging. In this study, we used GlycoSHIELD (GS), a powerful computational tool that allows us to generate 3D models of glycoprotein structures at a fraction of the computational time. GS generates glycoprotein trajectories where the protein remains static, and glycans explore all the conformations allowed by predefined criteria for steric clashes. We used CG criteria for determining clashes, where a glycan conformer is accepted if the distance between C-*α* of the protein and the glycan ring oxygens are above 3.5 Å. It is important to note that GS does not include glycan-glycan interactions. In the future, we plan to do MD simulations of glycosylated PNGs to study the effect of glycan-glycan interaction as well as the effect of protein flexibility on the structure of the glycoprotein, and their interactions with bNAbs. Despite these drawbacks, glycoprotein structures obtained using GS reproduce experimental structural features ^57^. GS thus allows faster analysis and understanding of the impact of glycans on protein structures and antibody-protein binding. We used Man9 glycans to model the HIV-1 glycoprotein structure in this work. GS prints separate trajectories for individual glycans.Chakraborty *et al*.^53^ observed minor variation in glycan topology for native-like vs Man9 glycosylation. To compare the impact of native-like glycosylation, we carried out all the analyses for clade A using the native glycans reported by Chakraborty *et al*.^53^. For each clade, we used N-glycosite^65^ predicted N-glycosylation sites for glycosylation. Figure 2(A) shows the PNGS for all the clades. Figure3(A) shows clade B structure with one Man9 glycan (as blue sticks) per site and Figure 3(B-D) display 160 glycan conformers per PNGS for clades A, B, and C. Trimer orientations are consistent with Figure 1.

**Figure 1.**
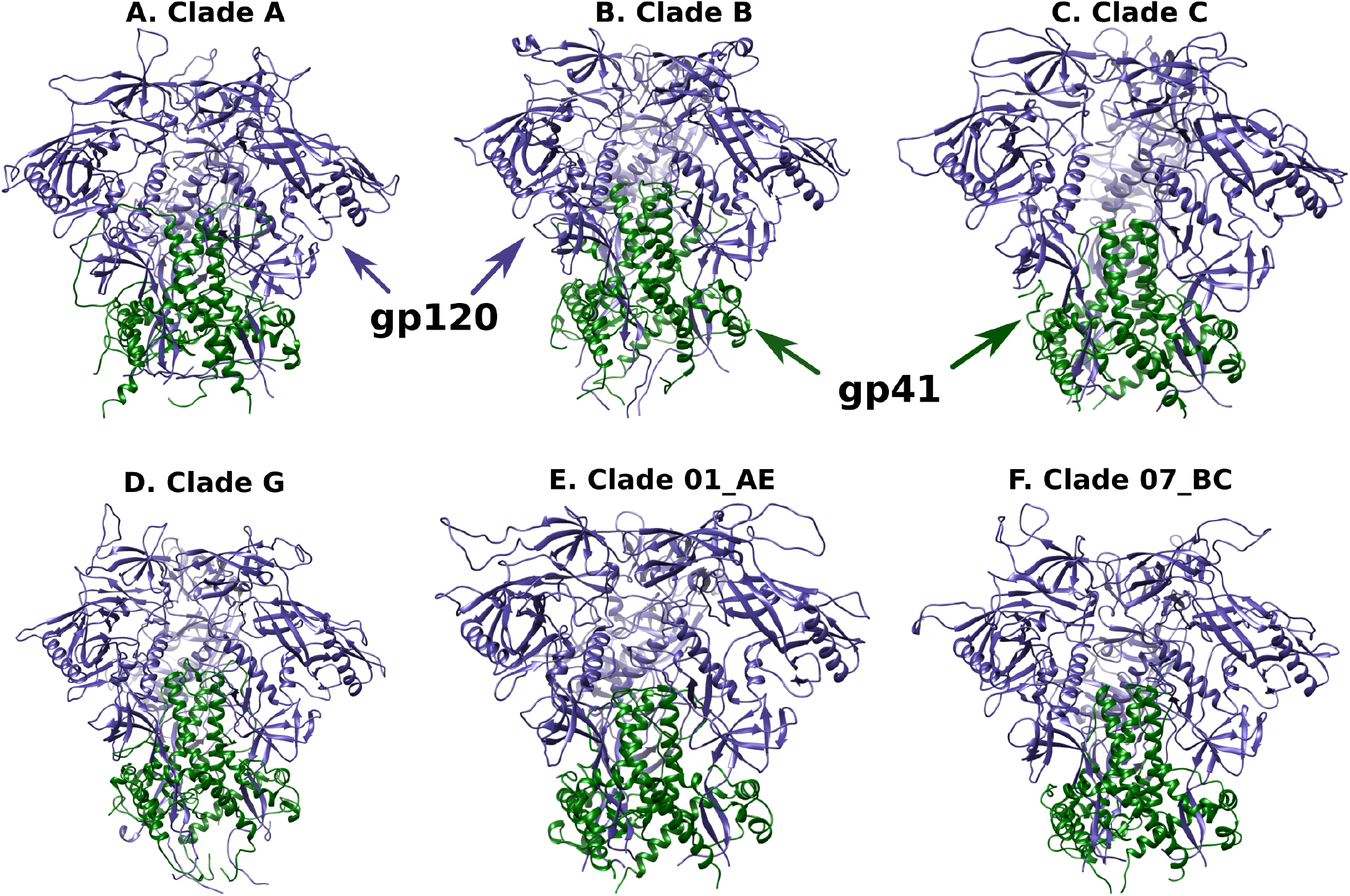
Prefusion crystallographic and cryo-EM structures of trimeric HIV-1 Env with modeled missing residues of clades (A) A (strain: BG505.T332N, PDB ID: 5FYL), (B) B (strain: JR-FL, PDB ID: 5FYK), (C) C (strain: CH848, PDB ID: 8SAU), (D) G (strain: X1193.c1, PDB ID: 5FYJ), (E) 01 AE ,(strain: BJOX018000.e12, PDBID 8GPI) and (F) 07 BC (strain: BJOX016000.e15 , PDBID 8GPJ).

**Figure 2.**
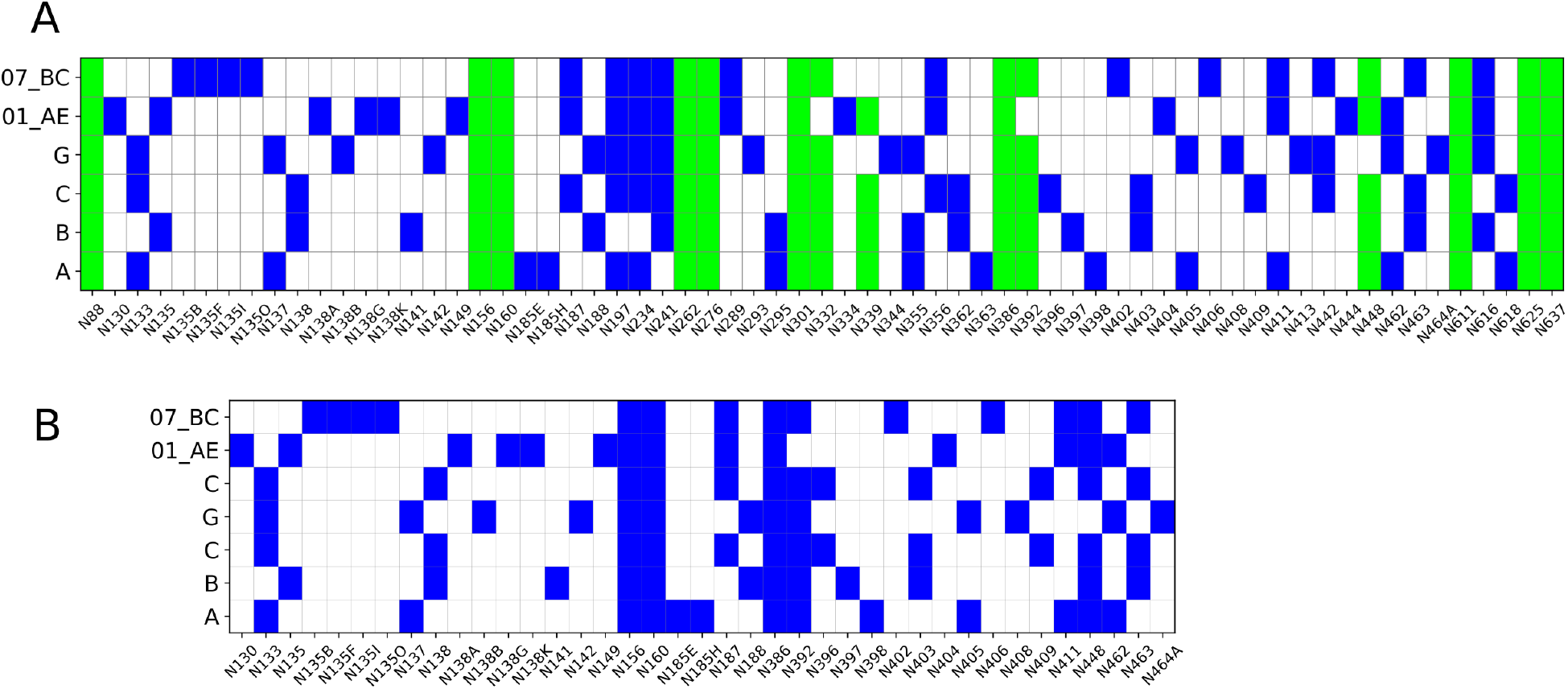
A) Potential N-glycosylation sites (PNGS) in each protomer for clades A (28), B (26), C (28), G (30), 01 AE (30), and 07 BC(29) obtained using N-glycosite program. AA sites are numbered based on the Hxb2 reference sequence. For a given clade, PNGS are shown in blue. PNGS conserved in clades A, B, and C are colored green. Note that native clade A (BG505) has no PNGS at site 332. B) PNGS considered for glycosylation in the set B structures are marked blue for each clade. All the selected PNGS are present in the flexible variable loop regions.

**Figure 3.**
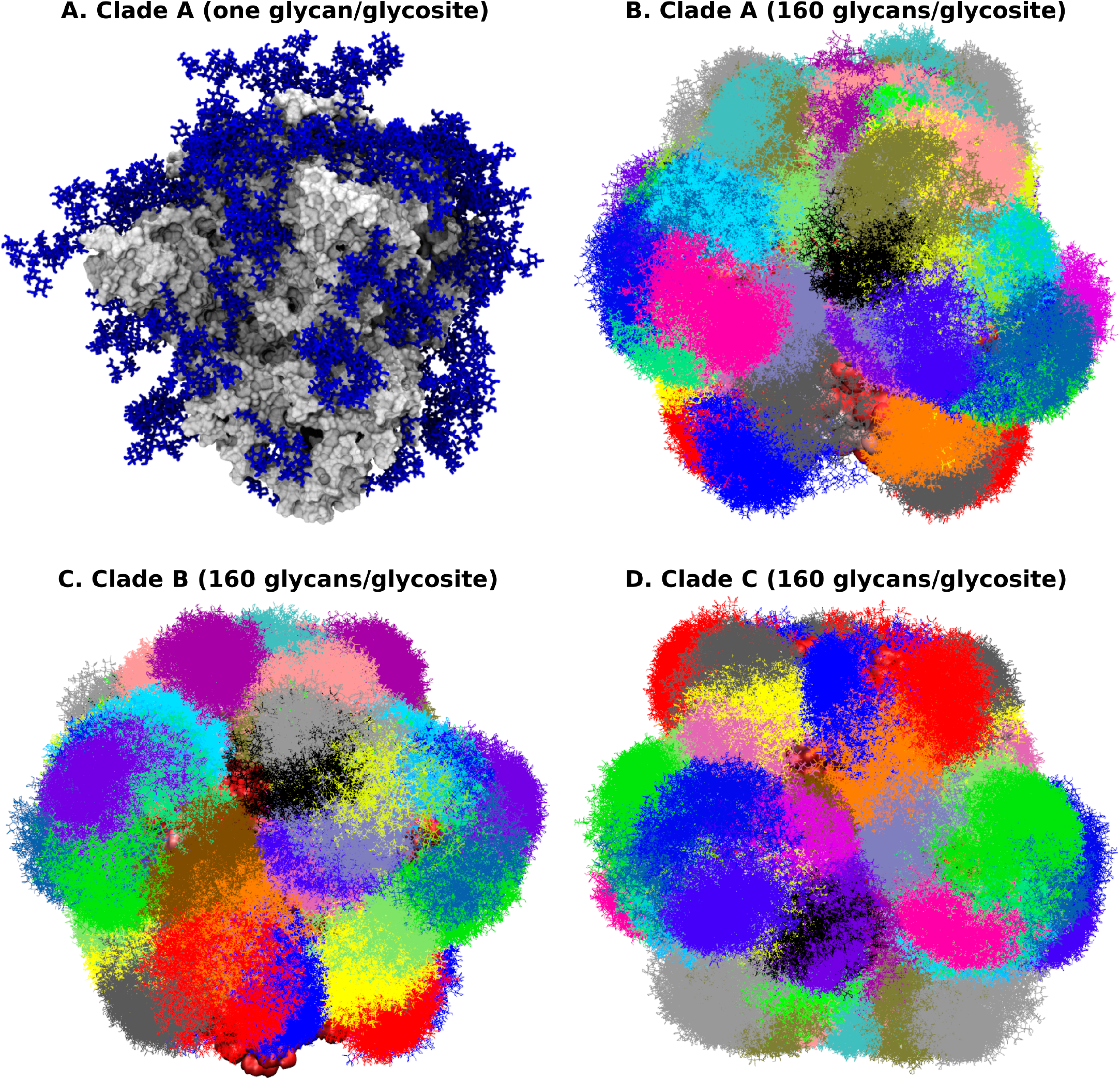
A) Glycosylated trimeric Env structure of clades B (shown as gray surface) with a single Man9 glycan conformer (shown in blue in stick representation) per glycosylation site. (B), (C) and (D) show glycosylated trimeric Env structures of clades A, B, and C with 160 Man9 conformers per glycosite. As shown, glycans shield the majority of the protein surface on average. As there are many overlapping regions, the removal of a particular glycan can be compensated by other glycans, maintaining the shield as observed in prior studies.

### Selection of protein structures

As the assumption of static protein structures may not work well for flexible loop regions, we performed short 10 ns long MD production of each clade and selected 11 geometries as described in the next section. These, along with each clade’s modeled crystal structure geometries, were chosen to generate the glycoprotein trajectories. Thus, we have a total of 12 protein structures for each clade. Figure 4 shows all 12 structures superimposed on each other for all clades. All the PNGS of the last selected frame from the MD simulation and the modeled crystal structure geometries (referred to hereafter as set A) were glycosylated using GS. However, only the PNGS in the flexible regions, see Figure 2(B), were glycosylated for the remaining 10 MD simulation frames (referred to hereafter as set B). For each glycosite, GS attempted to place 3000 glycan conformers for all the set A geometries and 500 glycan conformers for all the set B geometries.

**Figure 4.**
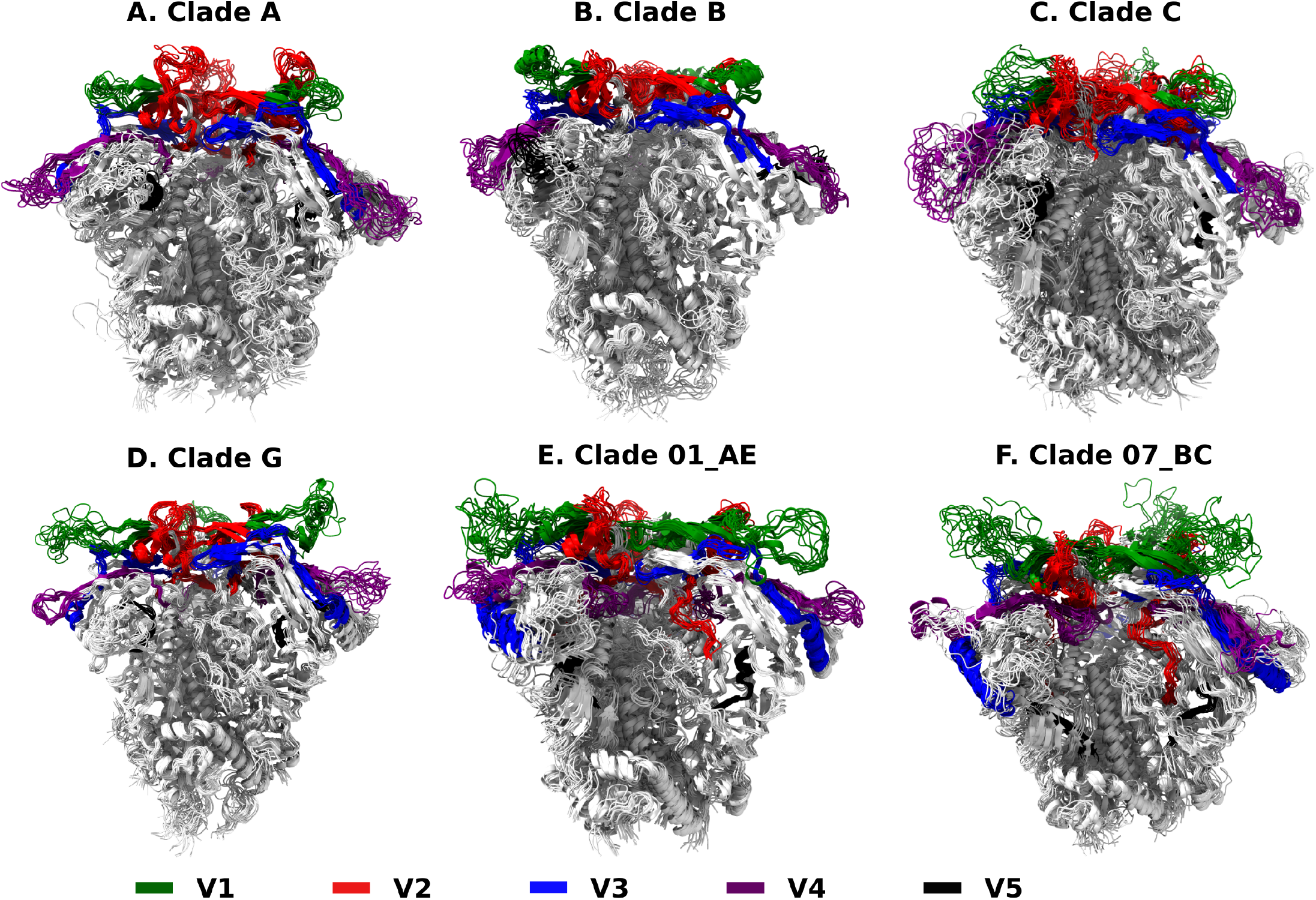
11 HIV-1 Env trimers obtained from MD simulations overlayed onto the crystal/cryo-EM structure of each of the clades (A) A, (B) B, (C) C, (D) G, (E) 01 AE, and (F) 07 BC. Variable loop regions V1 (green), V2 (red), V3 (blue), V4 (purple) and V5 (black) are shown. The cartoon representation of the remaining (non-variable regions) AA residues are shown in a light gray. For each clade, all the PNGS are used to generate glycoprotein trajectories for the crystal structure and the final structure from 10ns MD simulation (referred to as set A in the text). Whereas only the PNGS in the variable loop regions are used for the remaining 10 geometries (referred to as set B in the text). Figure 2 shows the PNGS used for glycan modeling for structures in sets A and B.

## MD Simulations

All the atomistic molecular dynamics simulations were performed in GROMACS v2022.3^66^ for the non-glycosylated trimeric Env of various clades. The PDB IDs of respective clades are 5FYL (clade A), 5FYK (clade B), 8SAU (clade C), 5FYJ (clade G), 8GPI (CRF01 AE), and 8GPI (CRF07 BC). After successfully modeling the missing residues (see sections above), we employed the CHARMM36 force field^67^ to generate the topology and associated coordinate files of the non-glycosylated HIV-1 trimer. The trimers were placed 1.5 nm from each side of the octahedron box. Subsequently, the boxes were solvated using the TIP3P water model.^68^ The overall charge neutrality in the systems was maintained using Na^+^ and Cl^*−*^ counter ions, and, additionally, 150 mM NaCl salt was added to the system. After this, we conducted 10000 steps of energy minimization to remove any bad contacts using the steepest descent method, and during this we kept heavy atoms of the trimers restrained using a restraint force of 500 kJ/(molnm^2^). This was succeeded by an additional 10000 steps without any restraint forces. We then performed NVT equilibration for a total of 500 ps, during which the heavy atoms of the trimer remained restrained at 500 kJ/(molnm^2^). The system temperature was maintained at 300 K using the Nose-Hoover coupling method^69,70^ with a coupling constant of 1.0 ps. The isotropic pressure in the system was ensured using the Parrinello-Rahman coupling method^71^ with a 5 ps coupling constant. The LINCS algorithm was used to constrain all the hydrogen bonds.^72^ The Verlet cutoff scheme was employed to calculate the non-bonded intermolecular interactions. Long-range electrostatic interactions were calculated using the Particle Mesh Ewald (PME) method.^73,74^ Finally, an NPT simulation was carried out without any restraints in the trimer for 10 ns, and trajectory frames were recorded every 5000 steps. A total of 11 frames of all the systems were extracted from the last 5 ns of the trajectory by skipping 40 frames between two successive extracted frames. These 11 coordinate frames were used for further analysis.

### Glycan-bNAb overlap analysis

To understand how bNAbs interact with HIV-1 Env glycans and how the interactions vary with clades, we calculate the extent of glycan occupancy within the binding volumes of bNAbs using the GS trajectories of the glycan-grafted HIV-1 trimers. We followed the procedure outlined by Stewart-Jones *et al*.^24^ for glycan-antibody overlap analysis. We superimposed the gp120 segment of the bNAb-HIV-1 to the corresponding segment on each HIV-1 gp120 monomer in the Env trimer of each clade. We extracted the coordinates of each bNAb obtained after the alignment and used them for the overlap analysis. As the protein structure remains static in GS, we align only to the first frame of each glycoprotein trajectory. We used the Matchmaker program in ChimeraX^75,76^for the alignment. Figure S3 provides a schematic description of the alignment procedure. Subsequently, we computed the average number of glycan atoms that are within 3.0 Å of the bNAb. 𝒩^*ij*^ denotes the number of overlaping atoms of the *i*^th^ glycan with the *j*^th^ monomer of the Env trimer. As each monomer may be present in different conformations during the MD simulation, 𝒩^*ij*^ can vary for each pair. For each bNAb and clade, three 𝒩^*ij*^ values are obtained corresponding to each of the aligned bNAb-monomer pairs. We renumbered the monomer chains before averaging the three 𝒩^*ij*^ values. The monomer to which the bNAb is bound is numbered 1, and the remaining two monomeric units are numbered 2 and 3 in the anticlockwise direction, consistent with the direction in the crystal structure. We denote this average over monomers as 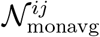. Next, we aver-aged 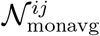 over all the 12 perfusion trimeric structures for each clade to obtain 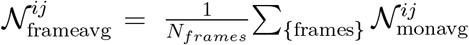. Where, *N*_*frames*_ = 12 for glycan sites considered in the set B struc-tures and *N*_*frames*_ = 2 for the remaining sites. We also computed the effective number of overlapping atoms, 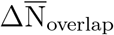, defined as 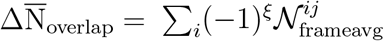, where *ξ* = 0(1) for glycans known to be important (unimportant) for bNAb’s recognition or binding.Although a few of the bNAb structures only have Fv region (shown in blue in Table S2), it should not affect our results as for most bNAbs the glycan atoms closer to the missing Fab region would also be within 3.0 Å of the Fv region.

### Total and relative accessible surface area

We computed the relative solvent accessible surface area (ΔSASA) (defined below) for each AA residue from the glycosylated trajectory of each clade using the GlycoSASA routine as implemented in GS. GS merges the individual glycan trajectories before computing the residuewise ΔSASA. Based on user input, it gives either the maximum ΔSASA or the average ΔSASA of each residue. Maximum ΔSASA at an AA residue is the maximum shielding due to each glycosylation site, and the average ΔSASA is the average shielding due to each glycosylation site. One drawback of merging the glycan trajectories is that the lowest number of accepted conformers, N_Acc_, of all glycosites limits the length of the merged trajectories. This might not be a problem if all glycosites have a similar number of accepted glycan conformers (N_Acc_). However, we observed that N_Acc_ varies with glycosites (see Figure 5) for all clades. We circumvent this problem by conducting the analysis separately for each glycan trajectory instead of the merged trajectory. Then, we combine all trajectory results to get the residuewise ΔSASA as described below.

**Figure 5.**
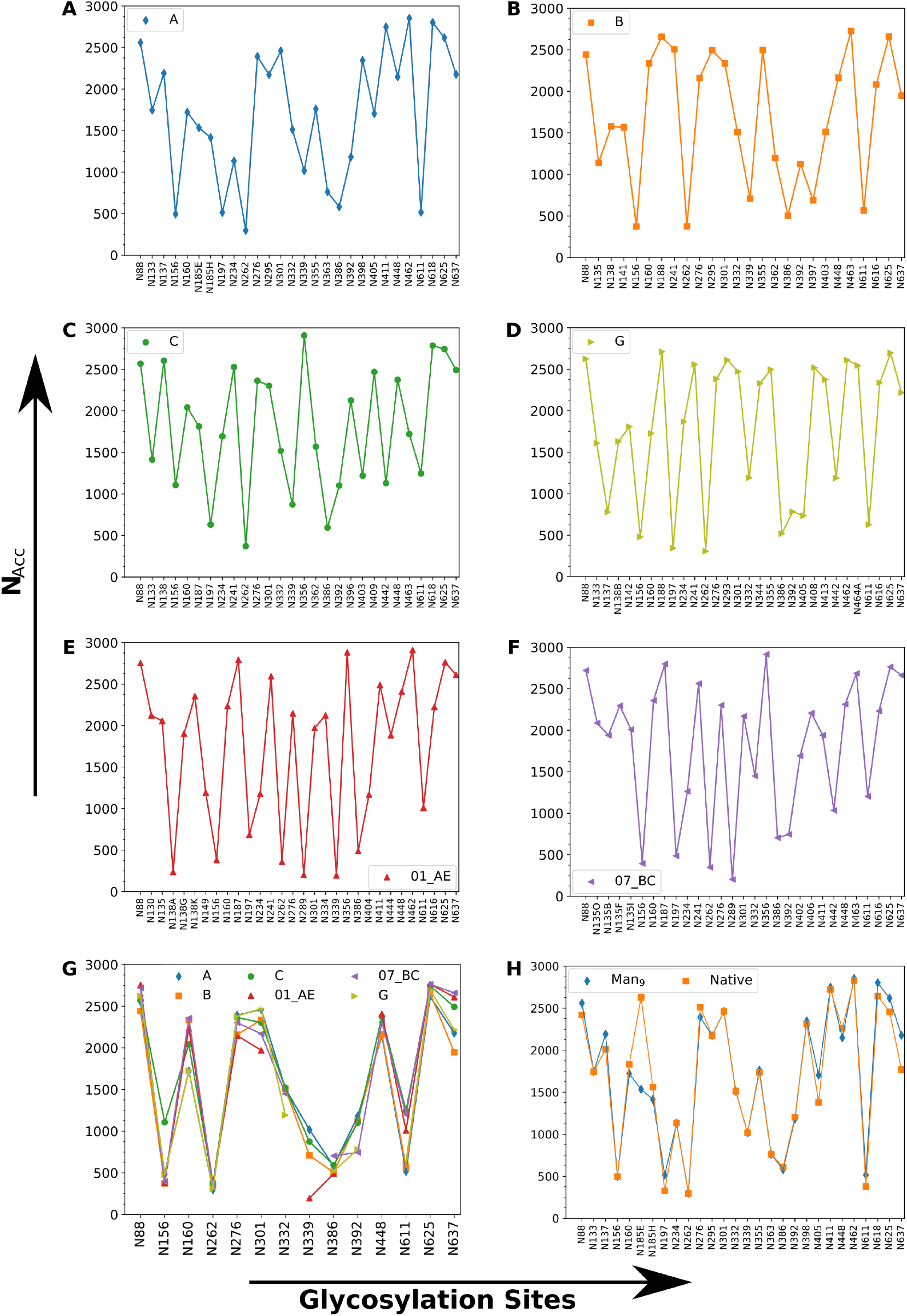
Variation of Number of accepted glycan conformers N_Acc_ per site out of 3000 conformers (averaged over all the 12 Env structures) obtained using GlycoSHIELD open-source software ^57^. (A-F) shows N_Acc_ for clades A, B, C, G, 01 AE, and 07 BC, respectively. Sites with more protein crowding around have a smaller N_Acc_ value. (G) N_Acc_ for conserved sites in clades A, B, and C. Barring a few sites, most glycosites exhibit clade-specific variations. (H) Comparison of N_Acc_ for Man9 and Native-like glycans for clade A. Except for site 185E, all other glycosite have similar N_Acc_, suggesting Man9 glycans can effectively model Native-like glycan patterns.

Relative solvent accessible surface area, ΔSASA, is defined as:

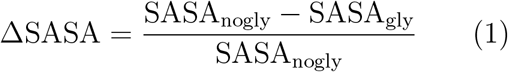

Here, SASA_nogly_ and SASA_gly_ are solvent-accessible surface areas for the bare and the glycosylated protein, respectively. A ΔSASA value of 1 for a given amino acid (AA) site indicates that the site is completely inaccessible when glycans are present.^57^ Conversely, a ΔSASA value of 0 means that the site is completely accessible even with glycans.^57^ Thus, a higher ΔSASA implies greater shielding. We calculated ΔSASA as follows. For each clade and static protein configuration,

1. Obtain the trajectory generated by GS for the trimer with a glycan attached at the *i*^th^ glycosylation site.
2. Using this trajectory, calculate ΔSASA at *j*^th^ AA site due to the above *i*^th^ glycan, denoted as 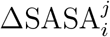, using Eq. 1.
3. Compute average ΔSASA^*j*^ at *j*^th^ AA site. This average is calculated by sum-ming the 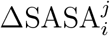 values across all gly-cosylation sites; i.e., 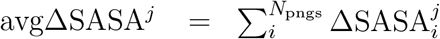. Additionaly, we computed the maximum ΔSASA^*j*^, maxΔSASA^*j*^, for the *j*^th^ AA site, as 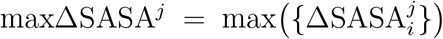. For protein structures in set B, 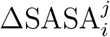 due to the absent glycans are taken to be the same as the corresponding value in the MD protein structure in set A.

Next, we computed average maxΔSASA^*j*^ and avgΔSASA^*j*^ over all frames. Moreover, we obtained SASA_gly_ from Eq. 1 by calculating the bare protein SASA of each residue, SASA_nogly_. We used Gromacs (Version 2019.6)^64^ for calculating SASA_nogly_.

### Epitope shielding calculations

We calculated the total glycan shielding of antibody epitopes, 𝒮, which quantifies the degree to which glycans shield the contact sites of bNAbs. For each bNAb, 𝒮 is defined as the sum of avgΔSASA over the epitope sites. , i.e. 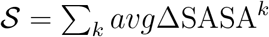, where *k* denotes the epitope sites. 𝒮 was then averaged over all the 12 protein structures per clade. We collected the contact site information for all the bNAbs from the LANL Neutralizing Antibody Contacts & Features repository^77^. A lower 𝒮 value might imply higher neutralization efficacy of the bNAb, thus helping estimate clade-specific efficacies of bNAbs.

## Results and Discussion

### Clade-specific variation in glycan shielding

#### Clade-specific variation in glycan conformers per site (N_Acc_)

Clade-specific variations in glycan shielding can significantly influence HIV-1-bNAb binding. To investigate these differences, we analyzed the number of glycan conformers (N_Acc_) successfully introduced by GlycoSHIELD (GS) at each glycosite across various HIV-1 clades. N_Acc_ serves as a qualitative measure of glycan conformational dynamics, with smaller values indicating less volume for glycan movement. Clade-specific differences in glycan shielding can result from variations in local protein structure or non-conserved glycosylation sites. Figure 2A indicates the potential N-glycosylation sites for different HIV-1 clades, as determined using the N-glycosite program^65^.

Figure 5(A-F) illustrates how the number of accepted glycan conformers (N_Acc_) across all potential N-glycosylation sites (PNGS) vary with clades A, B, C, G, 01 AE, and 07 B. Table S3 gives the corresponding values of N_Acc_. Our analysis reveals that N_Acc_ fluctuates significantly depending on the glycosylation site within each clade. While the majority of sites exhibit N_Acc_ values ranging from 1,000 to 2,500 out of 3,000 possible conformers (averaged over 12 trimer geometries), a smaller subset shows N_Acc_ values below 500, indicating increased steric clashes with nearby protein residues. Site N339 in clade 01 AE has the lowest number of accepted conformers with N_Acc_ = 196, while site N356 in clade 07 BC has the highest with N_Acc_ = 2,914.

Figure 5(G) compares the variation in N_Acc_ for conserved glycan sites among clades A, B, and C. At site N156, while clades A, B, and 01 AE show similar N_Acc_ values (300-450), clade C exhibits a significantly higher value (*>* 1000). This variation at N156, crucial for V1/V2 apex bNAb recognition (e.g., PG9, PG16), suggests potential differences in bNAb binding affinities across these clades. Similarly, N_Acc_ values for site N339 for clades A, B, and C are more than double that of clade 01 AE. Similar clade-specific variations are observed at glycosites N160 and N301 on the gp120 subunit, as well as N611 and N637 on gp41. Interestingly, site N332, known to be critical for many bNAb-HIV-1 interactions (e.g., PGT121, PGT128, PGT135, 10-1074) ^78^, shows no significant variation in N_Acc_ across clades

Figure 5H compares the number of accepted glycan conformers (N_Acc_) between native-like and Man9 glycans for clade A. We find minimal differences in N_Acc_ between these two glycosylation types across most sites, with the exception of site N185E. This finding aligns with previous research by Chakraborty *et al*.^53^, supporting the use of Man9 glycosylation as an effective model for native-like glycan patterns in HIV-1 studies.

#### Accessible surface area

The immunogenicity of HIV-1 is heavily influenced by the exposed surface areas on the Env protein^4,15,25,79^. Glycan holes, which are areas lacking glycan coverage, often direct the autologous immune response towards these regions. However, the resulting antibodies typically lack breadth, highlighting the importance of Env surface accessibility in vaccine design^25^. Furthermore, clade-specific variations in epitope accessibility may provide insights into antibody efficacy across different HIV-1 clades.

To investigate this, we used GlycoSHIELD (GS)-generated glycoprotein trajectories to compute the total and relative accessible surface area (ASA) of the Env trimer for six HIV-1 clades. We employed probe radii of 1.4 Å (corresponding to a water molecule), and 7.2 Å and 10 Å (representing single immunoglobulin domains, as estimated by Pancera *et. al*.^80^, Zhou *et. al*.,^25^ and Grant *et al*.^81^). Following the approach in Zhou *et. al*.^25^, we excluded residues within 15 Å of positions 31, 505, or 664 from ASA calculations for all clades. We included the N332 glycan for clade A, despite its absence in native clade strains, due to its presence in BG505-based immunogens used in vaccine design.^6,82^

Table 1 shows the variation in total accessible surface area (ASA) across HIV-1 clades with Man9 glycosylation, using probe radii of 1.4, 7.2, and 10 Å. Fully glycosylated and bare Env trimers of clades A and B show the highest solvent ASA, while clades G, 01 AE, and 07 BC show the lowest. Clade B exhibits the maximum immunoglobulin-domain ASA, whereas bare clade G has the minimum for both 7.2 and 10 Å probes. These findings suggest that clade B (JR-FL) may be the most immunogenic Env, while clade G (X1193.c1) the least. Glycosylation effects are consistent across clades, reducing solvent ASA by approximately 15% and immunoglobulin-domain ASA by about 55% (7.2 Å probe) and 62% (10 Å probe), respectively. Interestingly, ASA values for both glycosylated and bare Env show minimal variation between 7.2 Å and 10 Å probe radii, indicating that either could effectively mea-sure immunoglobulin-domain accessible surface area.

**Table 1.**
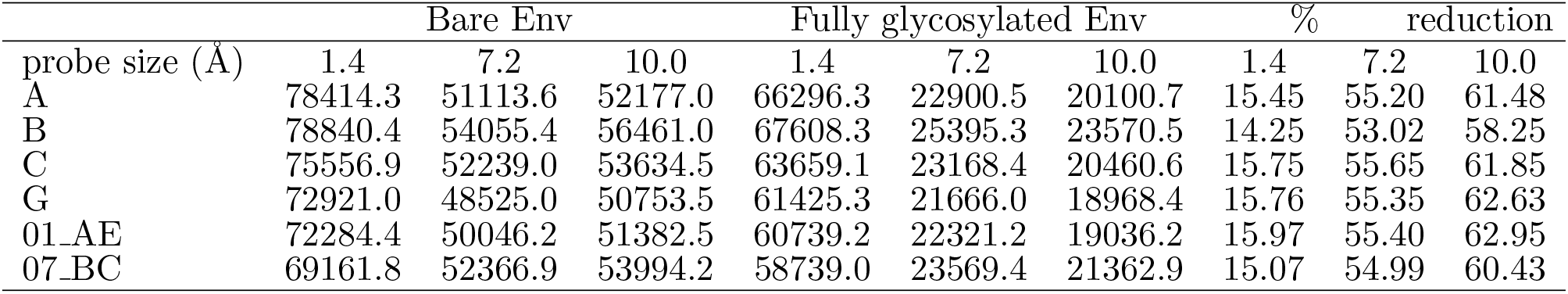
Total accessible surface area (Å^2^) for bare 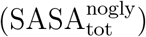 and fully glycosylated 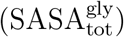 HIV-1 Envs of various clades with a probe radii of 1.4, 7.2, and 10.0 Å.

### Number of glycan contacts

All known bNAbs must navigate the glycan shield to bind to their epitopes ^10,24,55^. The number of glycan contacts can vary depending on both the specific bNAb and the HIV-1 clade. Following the approach of Stewart-Jones *et al*.,^24^ we quantified the number of contacts by counting glycan atoms within 3 Å of non-hydrogen bNAb atoms (see Methods section for details). Figure 6(A-C) presents histogram plots illustrating the number of bNAbs with an average of more than 10 (left) and 50 (right) glycan atom contacts per glycan, respectively. These plots are categorized by epitope classes and clades. Table S4 lists the corresponding numerical values. We note that our calculations do not account for the dynamics of bNAbs. Consequently, the number of glycan contacts observed in our analysis may differ from those in the actual Env-bNAb complex. Nevertheless, this analysis provides valuable insights into how bNAbs navigate the glycan shield and how this navigation varies across different clades. This information contributes to our understanding of bNAb-Env interactions and may inform future vaccine design strategies.

**Figure 6.**
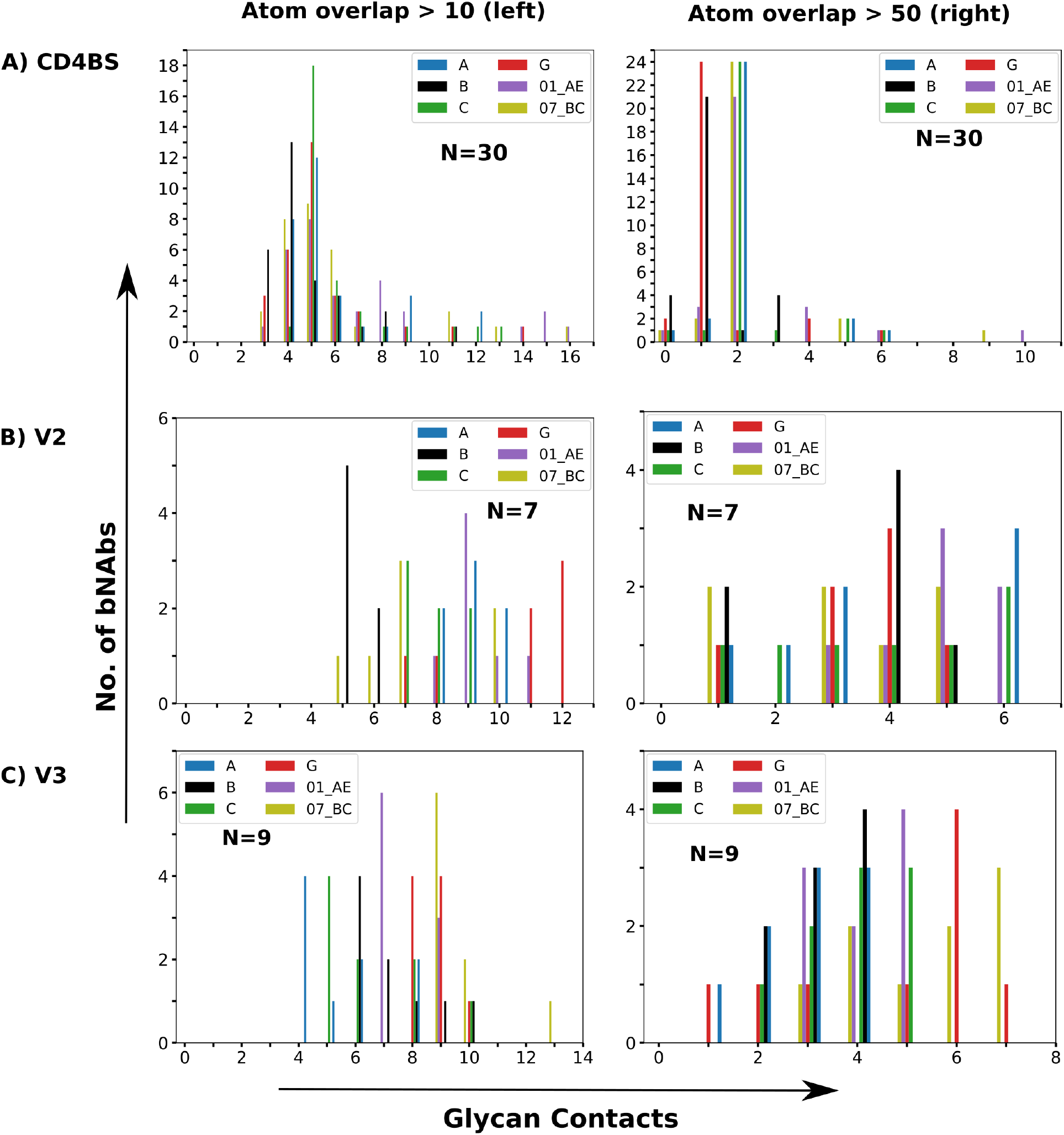
(Left) The number of Env glycans with at least ten glycan atoms on average occupying the bound volume of bNAbs, whereas (Right) plots the same with a threshold of 50 atoms. (A)CD4bs, (B) V1/V2 apex (V2), and V3-glycan targeting bNAbs. All bNAbs have an average of 10 glycan atom contacts with at least a single glycan.

Figure 6 illustrates substantial variation in glycan contacts among broadly neutralizing antibodies (bNAbs) within the same epitope class, even within a single clade. Out of the 30 CD4bs class bNAbs (see Figure 6A), we observed that clades B, G, 01 AE and 07 BC have at least one bNAb (clade B has six) interacting with glycans at three PNGS each making more than 10 atom contacts, the minimum. bNAb b12 interacts with glycans at 16 PNGS of clades 01 AE and 07 BC, each making more than 10 atom contacts, the maximum in this class. Most CD4bs bNAbs across all clades typically engage with 4-5 glycans, each making more than 10 atom contacts. Interestingly, all clades feature at least one CD4bs bNAb that makes no glycan contacts involving more than 50 glycan atoms. In clades B and G, the majority of bNAbs interact with one glycan making more than 50 atom contacts, while in the remaining clades, most bNAbs interact with two such glycans. Notably, in clade 01 AE, bNAb b12 engages with glycans at 10 PNGS, each making more than 50 atom contacts, which is the highest observed in the CD4bs class.

For V1/V2 apex bNAbs, as illustrated in Figure 6B, all antibodies interact with glycans at atleast five PNGS, each making more than 10 atom contacts. Clade B has none of its bNAbs in this class interacting with more than seven glycans at this contact level (more than contacts). At least 9 glycans of Clades 07 BC have more than 10 atom contacts, Only clades A, G, 01 AE, and 07 BC feature bNAbs that engage with glycans at more than 10 PNGS, each making over 10 atom contacts. while clades 01 AE and G has at least 7 and 8 glycans making more than 10 atoms contacts, respectively. All bNAbs in this class engage with glycans at more than two PNGS of 01 AE, each making more than 50 atom contacts, the highest. Only clades A, C, and 01 E has bNAbs that interacts with glycans at six PNGS, each making at least 50 atom contacts.

For V3-glycan bNAbs, as shown in Figure 6C, all antibodies interact with glycans at more than three PNGS, each making at least 10 atom contacts. At least 9 glycans of Clades 07 BC have more than 10 atom contacts, while clades 01 AE and G has at least 7 and 8 glycans making more than 10 atoms contacts, respectively. Only clades B, C, G, and 07 BC feature bNAbs in this class that engage with glycans at more than 10 PNGS, each making over 10 atom contacts. All clades have at least one bNAb that makes glycan contacts involving more than 50 atoms. For clades G and 07 BC, most bNAbs have cglycans at more than 4 PNGS, each making at least 50 atom contacts.

These findings highlight the complex and diverse nature of bNAb-glycan interactions across different HIV-1 clades. In the next section, we analyzed the Env glycan-bNAb interaction in extensive details.

### Glycan-antibody overlap analysis

#### CD4bs bNAb-glycan overlap

Figure 7 and 8 illustrate the glycan-bNAb contacts for CD4bs bNAbs CD4bs targeting bNAbs are usually associated with the best breadth and potency combination^83,84^. bNAbs in the CD4bs class are classified into VH1-2derived (e.g. VRC01-like Abs), VH1-46-derived (e.g. 1B2530 and 8ANC131) and antibodies derived from diverse VH-genes(e.g: b12, HJ16, CH103) ^85^. VRC01-like antibodies mimic CD4 binding to gp120, whereas antibodies derived from diverse VH-gene utilize a loop-based recognition mechanism^85^. VH1-46-derived antibodies generally have similar recognition mechanisms as VRC01-like antibodies^85^.

**Figure 7.**
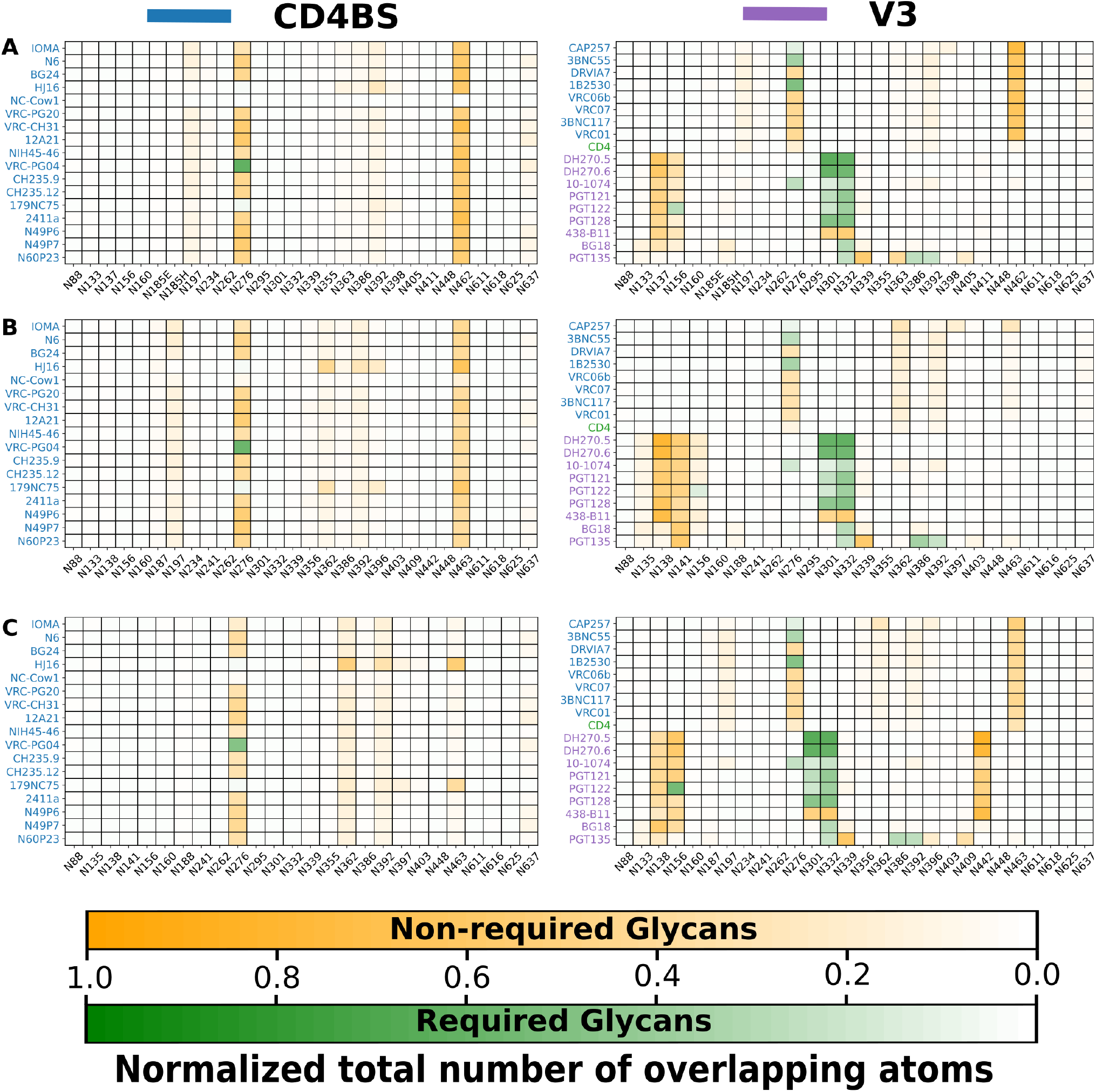
Glycan-bNAb overlap for CD4bs and V3-glycan bNAbs for clades (A) A, (B) B, and (C). Sites for which glycans are required for recognition by a bNAb are colored green, whereas glycans not known to be required for recognition are colored orange. Magnitude of 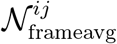 (see methods) determines the color intensity.

**Figure 8.**
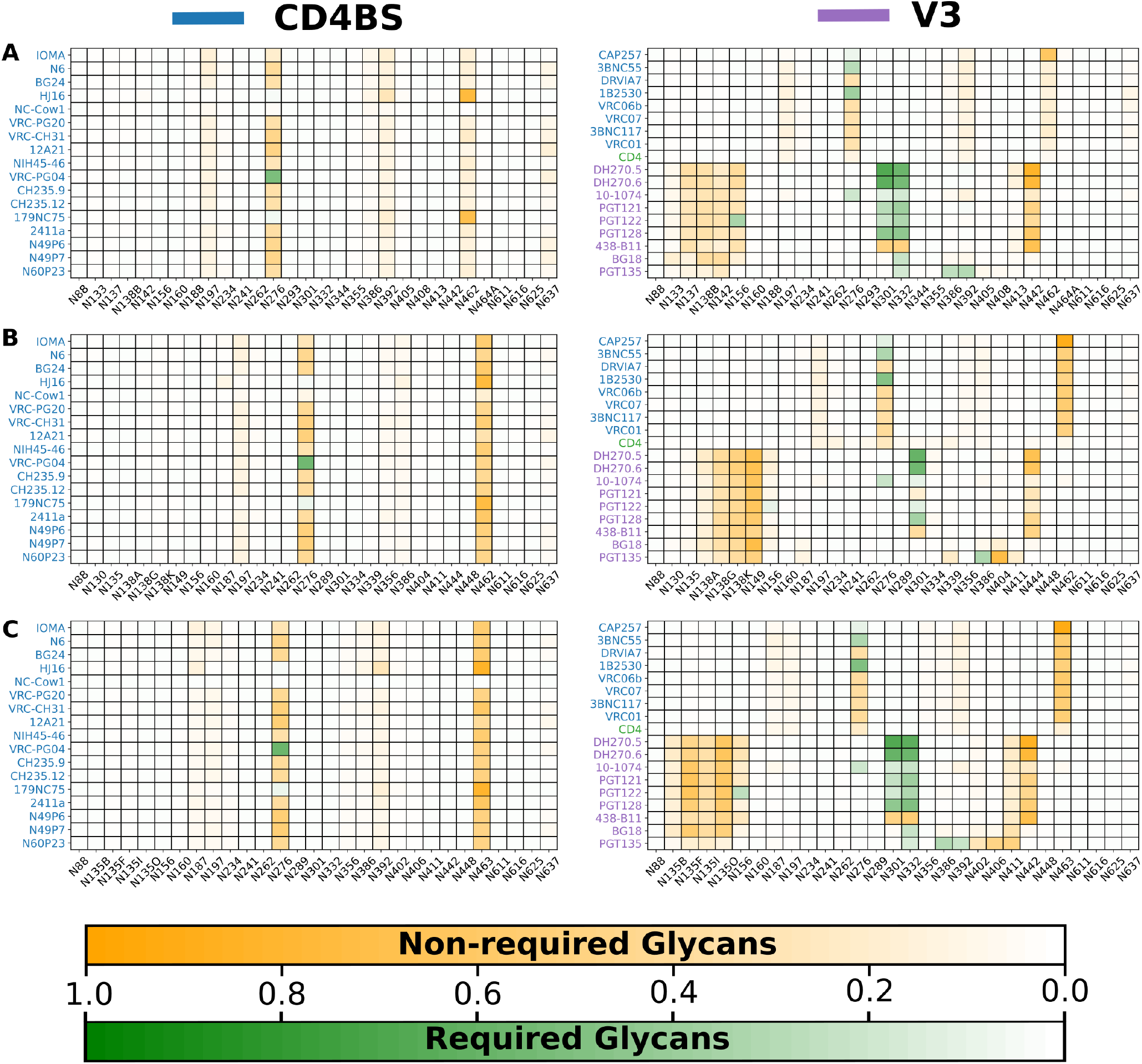
Glycan-bNAb overlap for CD4bs and V3-glycan bNAbs for clades (A) G, (B) 01 AE, and (C)07 BC. Sites for which glycans are required for recognition by a bNAb are colored green, whereas glycans not known to be required for recognition are colored orange. Magnitude of 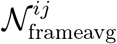 (see methods) determines the color intensity.

We first focus our attention on VRC01 and 3BNC117, two CD4bs bNAb that are known for their breadth and are extensively studied in clinical trials for HIV-1 treatment and prevention^86^. A phase 2 trial based on the long-acting form, 3BNC117-LS, is currently recruiting participants (trial identifier NCT06205602). As shown in Figure 7A, both VRC01 and 3BNC117 have similar glycan overlap patterns for clade A (BG505+N332). Both have substantial overlap with glycans N276 and V5 loop glycan N462 and, to a lesser extent, with the glycans at N197 and N392. In addition, we observed a minor overlap with the glycans at N234, N363, N386, and N637. However, overlap with N462 is significantly more than observed in ref. ^24^ for VRC01. While Stewart-Jones *et. al*.^24^ find no overlap with N234, N386, N392, and N637. The observed differences may be due to greater conformational sampling achieved by GS, which is harder to achieve in conventional MD simulations. Interestingly, Stewart-Jones *et. al*.^24^ observed significant overlap of VRC01 with N301 and the overlap with the glycan at N363 was more than what we observed.

Both VRC01 and 3BNC117 have similar glycan overlap patterns for all the clades except for clades B and G. The overlap with the glycan at site N463 in clade B (JR-FL) and at N462 in clade G (x1193c?1) is considerably less than other clades. Steric hindrance due to V5 loop glycans is a well-known contribut-ing factor to the resistance of many VRC01class antibodies^13^. Bricault *et al*.^84^ also observed a high number of PNGS in the V5 loop (N460/N462/N463) to be associated with higher resistance of CD4bs bNAbs. Minimal contacts with the V5 glycan site at N463 should favor greater binding of CD4bs bNAbs (VRC01, 3BNC117, N6, VRC07, N49P7, CH235.12) to clade B viruses compared to other clades.

To understand the difference in glycan contact at the site N462/N463 at the molecular level, we examined the glycan overlap with the VRC01 bNAb by visualizing the glycan-grafted GlycoSHIELD (GS) trajectories of clades A and B superimposed with the aligned VRC01 structure (see Figure 9). Figures 9 (A) and (C) present side views, while Figures 9 (B) and (D) show top views, displaying a total of 160 overlaid glycan conformers. We observed that glycan conformers are oriented toward and away from VRC01 for clades A and B, respectively. Figure 9 (E) shows superimposed structures of clade A (in blue), clade B (in green), and clade C (in red). These differences in glycan orientation are due to orientational differences of the N462/N463 sites in these clades.

**Figure 9.**
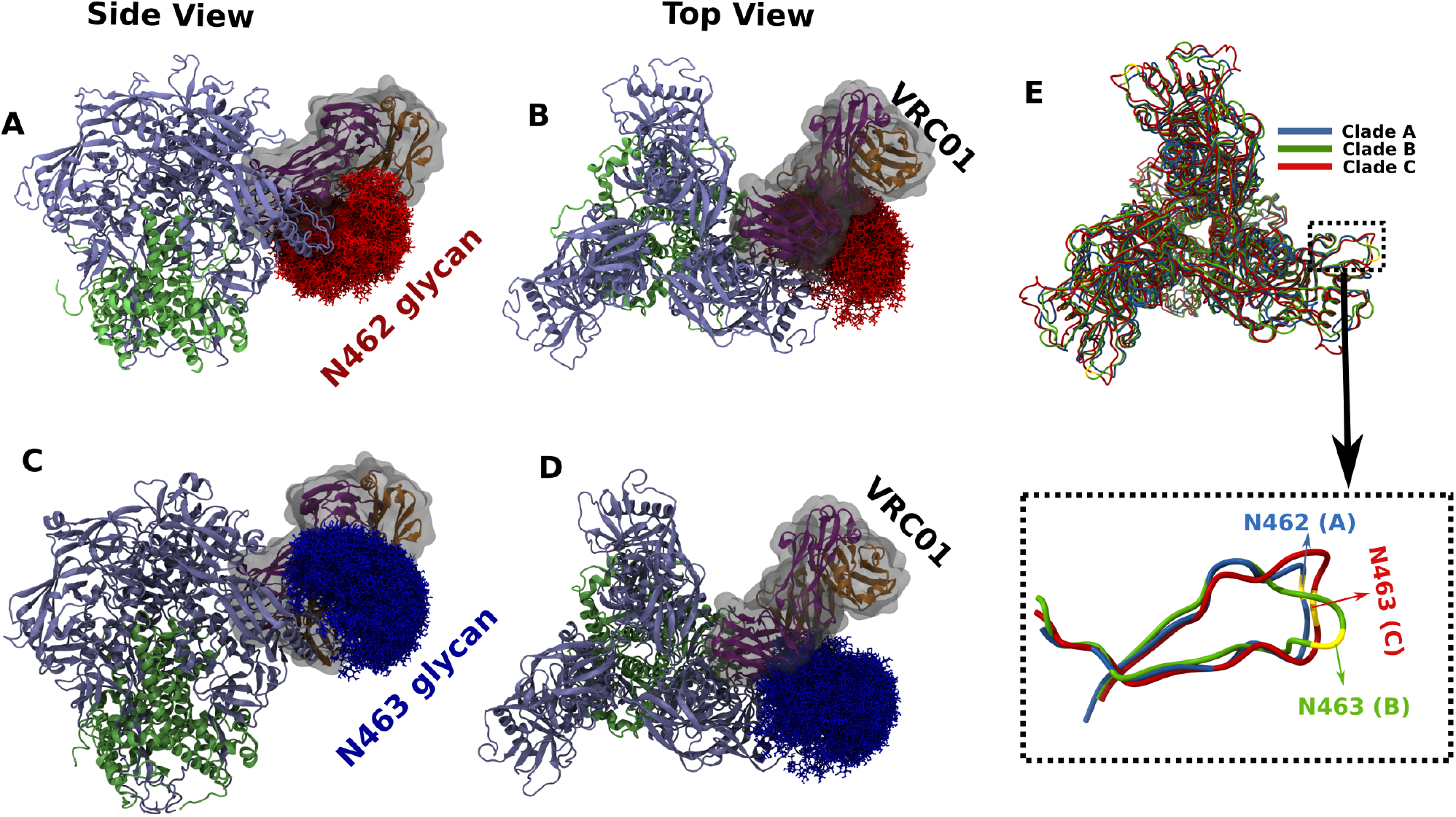
VRC01 interaction with glycan at N462/N463. VRC01 (gray surface, purple: heavy chain, orange: light chain) bound to clade A [(A) side and (B) top], and clade B [(C) side and (D) top] Trimers. Gp120 and gp41 are represented in iceblue and green color, respectively. Clade A glycans at N462 are shown in red (in the sticks), whereas clade B glycans at N463 are shown in blue (in the sticks). A total of 160 glycan conformers are displayed for each clade. Glycan and VRC01 have severe clashes with clade A N462 glycans [(A) and (B)]. In contrast, there is only a minor overlap of VRC01 with clade B glycans at N463 [(C) and (D)]. The orientation of glycans is distinct for the two clades. (E) On the left are superimposed structures of clades A (dark blue), B (green), and C (red). The magnified region of the AA residues surrounding N462/N463 is shown on the right. N462/N463 sites are shown in yellow color. The orientation of site N463 in clade B [red] is different from site N462 in clade A [green](right panel). These differences lead to the differences in the orientation of glycan conformers.

Next, we compare these bNAbs with other bNAbs in the VRC01-class; see Figure 7 and Figure 8. Within a clade, all the VRC01-like bNAbs have qualitatively similar glycan contacts. Thus, all the above analyses are expeted to be valid for the clades.

Compared to VRC01-like bNAbs, VH-gene-derived bNAbs HJ16 and NC-Cow1 have comparatively much less contact with N276 in all the clades. In addition, NC-Cow1 has a significantly lesser overlap with the glycan at N462/N463 compared to VRC01-like Abs in the same clade.

Figure 10(A) compares the mode of binding of VRC01 (dark blue) and HJ16 (red) to clade A structure. As shown in Figure 10A, all the bNAbs have different binding orientations.

**Figure 10.**
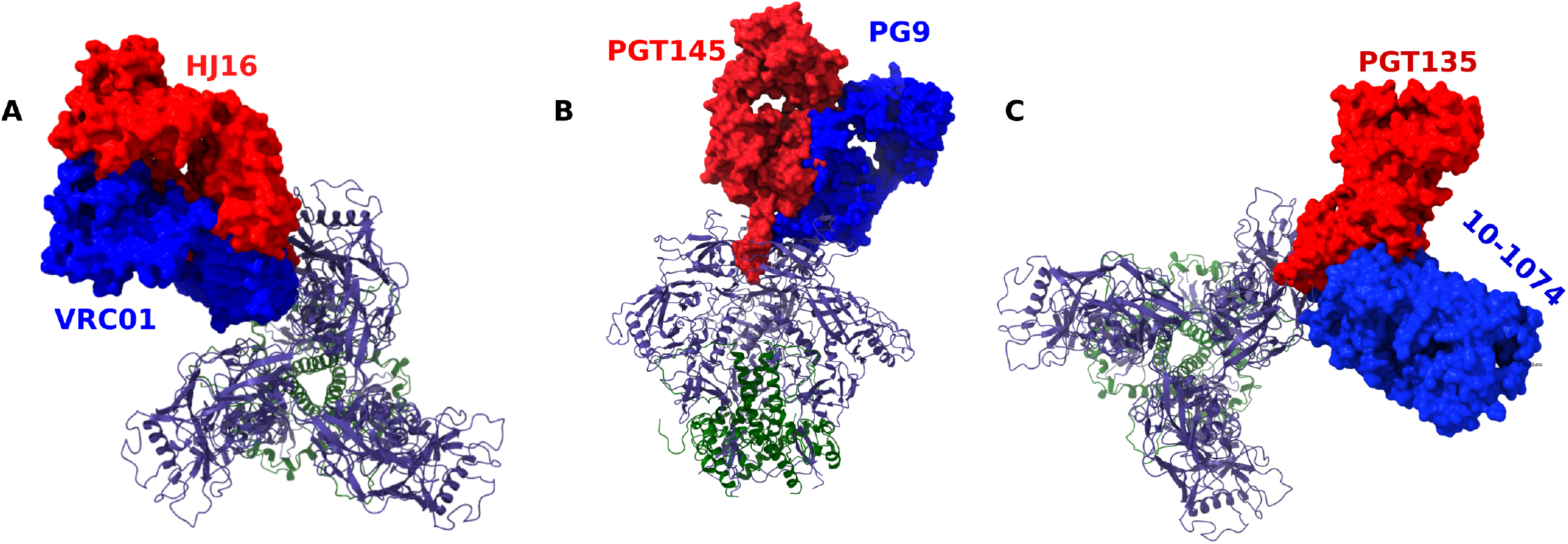
bNAbs approach in different angles for binding. Superimposed Clade A structure in complex with bNAbs (a) VRC01 (dark blue) and HJ16 (red), (b) PG9 (dark blue) and PGT145 (red), and (c) 10-1074 (dark blue) and (red). Gp120 and gp41 are shown in ice blue and dark green color. Env Trimers (gp120 :ice blue cartoon, gp41: green cartoon) are shown in top orientations in (a) and (c) and side orientation in (b) for visual clarity. Orientation differences in binding mode lead to different glycan interactions for bNAbs in a particular class. Note that, as the structure of PGT145 (PDB ID: 6NIJ) and 10-1074 (PDB ID: 7UCG) used in our analysis contain only the Fv region, we have used aligned full fab apo x-ray structures with PDB ID 3U1S for PGT145 and 4FQ2 for 10-1074

N6, a CD4bs bNAb, has high potency and breadth^79^. Its long-acting form, N6LS, has shown promising results in recent clinical trials^87^. N6 is known to avoid glycan clashes by shifting the orientation of attack^13,79^. However, we do not observe any substantial differences in glycan contacts of N6 from other VRC01class antibodies. This suggests that N6 has to navigate through a similar glycan barrier before binding to Env. Once bound, contacts might be reduced. Stewart-Jones *et. al*.^24^ observed similar discrepencies between simulation and experiments for VRC01.

IOMA, another antibody in the VRC01class, lacks some features that are standard to VRC01-class Abs. For example, it lacks the short CDRL3 that defines VRC01-class bNAbs and it requires fewer somatic hypermutations (SHM)^51,88^. However, we do not see changes in the overlap pattern between IOMA and other bNAbs in CD4bs. Gristick *et. al*.^51^, using high-resolution structures of IOMA in complex with HIV-1 Env, observed that some features of IOMA allow it to evade glycan shields and make it a good candidate for vaccine design. These results suggest that although a similar number of glycan atoms occupies the binding volume of antibodies, bNAbs can have different ways of interacting with or accommodating glycans once bound. While most CD4bs bNAbs are shielded substantially by N462/463 glycan, none is known to bind or accommodate this glycan. Hence, it might be beneficial to design an immunogen lacking the glycan at site N462.

#### V1/V2 apex bNAb-glycan overlap

bNAbs in the V1/V2 apex class target the trimer apex, composed of V1 and V2 loops. Figure 11 shows the glycan overlap patterns of the bNAbs in this class. Due to the quaternary nature of the epitopes of these bNAbs, they overlap with glycans from more than one Env protomer. As no bNAbs in this class have overlap with glycan sites with the Hxb2 site number greater than 300, only sites with number smaller than 300 of each protomer are shown. All of them are known to require N160 glycan for neutraliztion^89–91^, while N156 glycan is also stated to be necessary in a strain-specific way^92^. In clade A, PG9 has strong primary overlap with glycans at N137, N156, and N160 from one protomer and, in addition, has interactions with glycans at N160, N185E, 185H, and N197 from an adjacent gp120 protomer. PG9 interactions with N137, N185E, 185H, and N197 were not observed in prior experimental studies^93^. Although interactions with N137 were observed in computational studies^24^, they considered glycan contacts with a single protomer. PG9, hence, must navigate through additional glycans than previously thought. PG9 has a similar glycan overlap with all other clades, and the only variation comes from the additional or lack of glycans in the V1 and V2 loops. For example, all three V1 loop glycans at N130, N135, and N138A in clade 01 AE substantially overlap with PG9. However, 07 BC glycans at sites N135F, N135I, and N135O have only minor contact with PG9. For a given clade, most V1/V2 apex bNAbs, such as J033, J038, VRC26.25, VRC38.01, and PG16, overall have similar glycan overlap patterns as PG9. For the HIV-1 clades analyzed, PGT145 interact with glycans on all protomers of the trimer, though contacts with the third protomer are relatively weak. Contrary to the other bNAbs in this epitope class, PGT145 has no contact with the glycan at N156.

**Figure 11.**
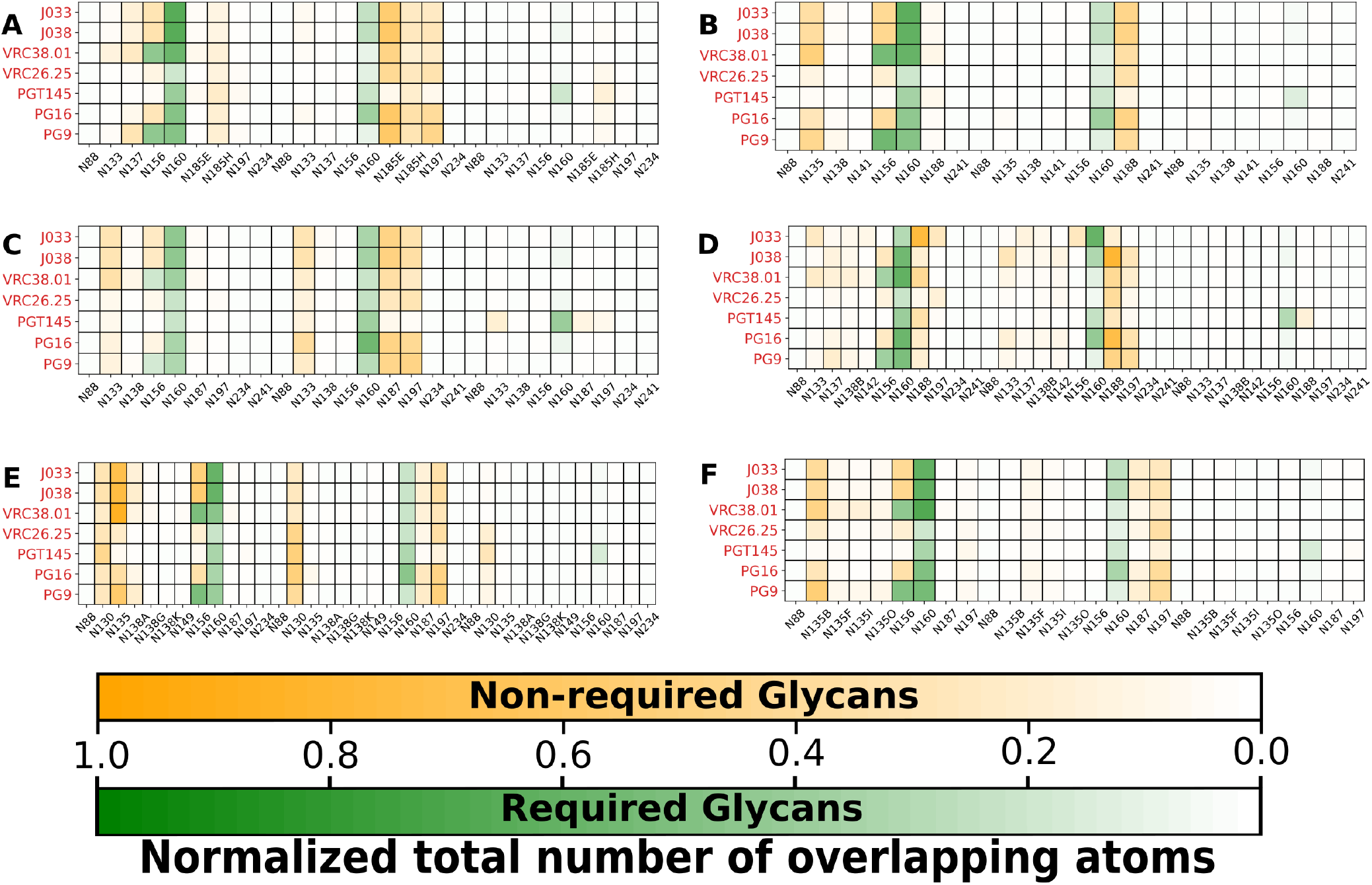
Glycan-bNAb overlap for V1/V2 apex bNAbs for clades (A) A, (B) B, (C) C, (D) G, (E) 01 AE and (F)07 BC. Sites for which glycans are required for recognition by a bNAb are colored green, whereas glycans not known to be required for recognition are colored orange. Magnitude of 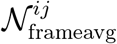 determines the color intensity. Only the glycan at sites smaller than 300 for each of the three protomers is plotted. Overlap with the absent glycan sites are zero.

This variation is due to the differences in the binding orientations of PG9,and PGT145. Figure 10B shows the Clade A structure in complex with PG9 (dark blue) and PGT145 (red). PGT145 appears to approach the trimer perpendicular to the V1/V2 apex, whereas PG9 attacks it at a tilted angle and is oriented away from the third protomer.

Removal of N197 glycan reduces the sensitivity or neutralization of PG9^4^ in clade A variant. Behrens *et. al* attributed this effect to glycan-glycan interactions whereby removal of N197 glycan perturbs the glycan at the trimer apex^4^. We see that for all clades PG9 has substantial overlap with N197 glycan, suggesting that N197 might directly affect PG9 binding. Although a similar reduction was observed for PGT145, we do not see interactions with the N197 glycan.

Overall, our findings reveal that V1/V2 apex bNAbs can have clade-specific differences in binding coming from glycans at the V1 and the V2 loops as well as from their variation in the approach for binding.

#### V3-glycan bNAb-glycan overlap

V3-glycan class bNAbs recognize the base of the V3 loop and associated glycans. These bNAbs are known to strongly depend on glycans at positions N301 and N332 for recognition ^10,11^. Figures 7 and 7(B) (right panel) compares the glycan-bNAb interactions for V3-glycan class bNAbs for various clades. V3-glycan bNAbs 10-1074 and PGT121 have been investigated in human clinical trials and have shown promise for HIV prevention and treatment. BG18 is also vital from a vaccine standpoint owing to the absence of insertion-deletion (indels) and its greater potency and similar breadth to other V3 targeting bNAbs^52,79,94^.

Except for PGT135 and BG18, all V3-class bNAbs considered here (10-1074, PGT121, PGT122, PGT128, 438-B11, DH270.5, and DH270.6) have similar glycan contacts within a particular clade.

In addition to the glycans at N301 and N332, all of 10-1074, PGT121, PGT122, PGT128, 438-B11, DH270.5, and DH270.6 interact strongly with V1 loop glycans. As clades G, 01 AE, and 07 BC have a higher number of V1 loop glycans compared to clades A, B, and C, V3-glycan bNAbs are expected to face greater glycan barriers, thereby affecting their activity. Notably, V4 glycans at N442/N444, which are absent in clades A and B, substantially overlap with these bNAbs.

The bNAbs BG18 and PGT135 lack contact with N301 and N442/N444 in all clades. In addition, PGT135 has significant contact with V4 loop glycans at N386, N392, N404/N406 compared to the others in this epitope class. The difference between PGT135 and other bNAbs in V3 glycans can be attributed to the binding orientation difference as shown in Figure 10C.

#### Fusion peptide, gp120-gp41 interface bNAb-glycan overlap

bNAbs in the gp120/gp41 interface class target a region spanning both the gp120 and the gp41 subunits^95–98^. Figure 12) reveals that 8ANC195 has contacts with several glycans on both gp120 (sites N234, N276) and gp41 (sites N616/N618, N637) in all clades, as shown in experiments ^99^.

**Figure 12.**
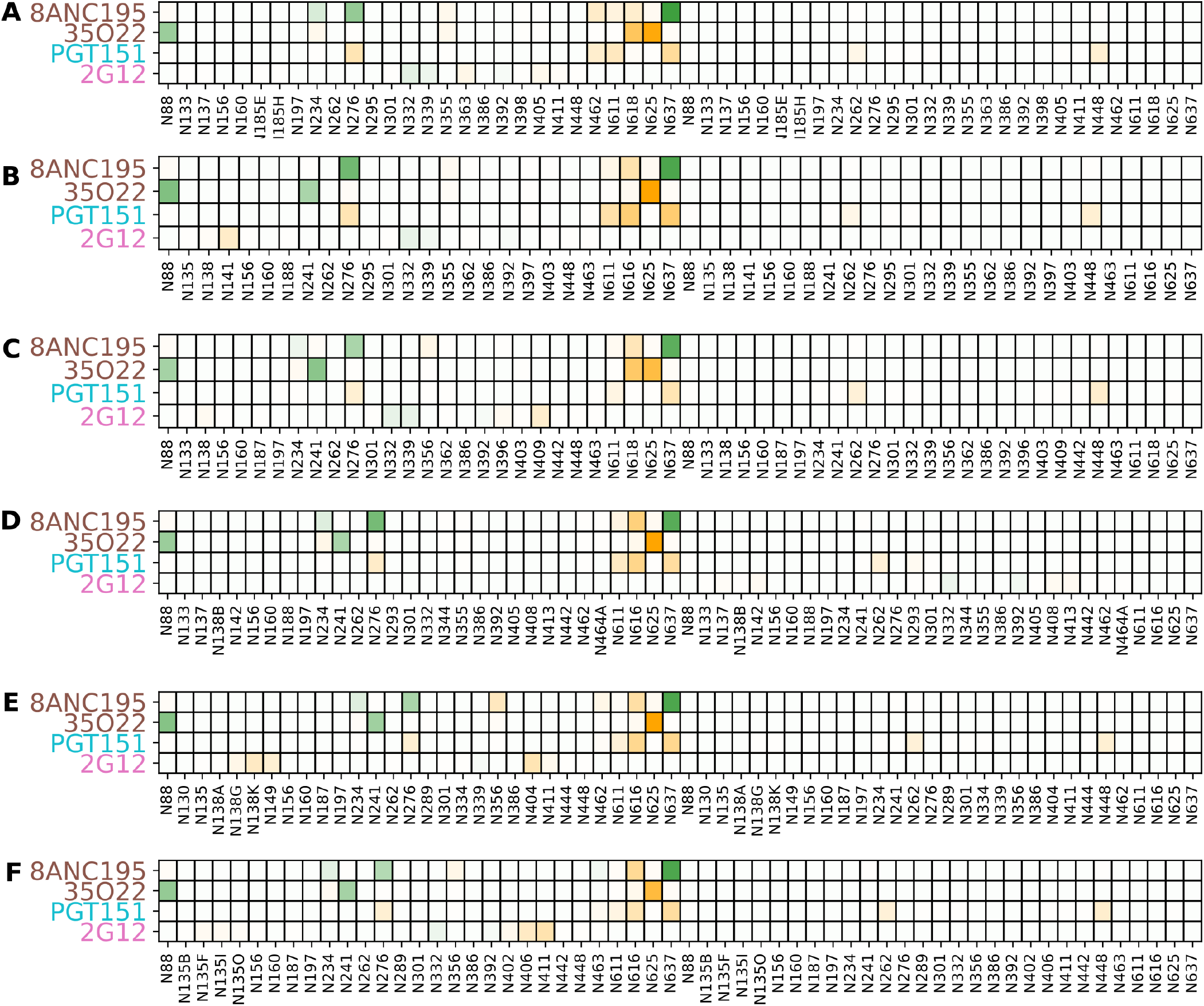
Glycan-bNAb overlap for bNAbs targeting gp120/gp41 interface, fusion peptide, and glycan for clades (A) A, (B) B, (C) C, (D) G, (E) 01 AE, and (F)07 BC. Sites for which glycans are required for recognition by a bNAb are colored green, whereas glycans not known to be required for recognition are colored orange. The magnitude of 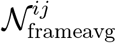 (see methods) determines the color intensity.

In addition, weaker overlap with glycans at sites N355/N356 and N611 was observed. Although weak, overlap with N462 glycan is observed in Clade A and 01 AE, which is missing with glycan at N463 in Clades B, C, G, and 07 BC.

35O22, another gp120/gp41 interface class bNAb, has different glycan overlap in all the clades from that of 8ANC195^100,101^. 35O22 primarily has overlap with glycans at sites N88, N241, N616 and N625 (see Figure 12). Clade A strain lacks N241, a glycan known to be required for 35O22 binding. In addition, it has a strong overlap with glycans at site N618 in clades A and C. No overlap with glycan at N618 in clades B, G, 01 AE, and 07 BC is observed, suggesting that these clades will have fewer obstructions for 35O22 binding compared to clades A and C.

The bNAb PGT151 targets the fusion peptide region of the HIV-1 Env.^50,83,102,103^ As shown in Figure 12, PGT151 interacts with PNGS at N611 , N616 , N637 , N276 of the one protomer of clade B, and with N262 and N448 of the second protomer. Figure S5 gives a pictorial depiction of PGT151 interactions with clade B glycans. Our results are overall consistent with the experiment of Lee JH *et al*.^104^. While Lee JH *et al*. predicted more interactions of PGT151 with clade B glycans at N448 and N241 compared to N276, we find more N276 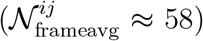 gly-can atoms inside the bound volume compared to both N448 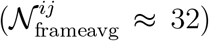 and 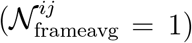. This suggest that PGT151 evades the N276 glycan to bind to the HIV-1. As observed in the experiments by Lee JH *et al*.^104^, glycans at N241 are in close proximity to PGT151(see Figure S5).

#### 2G12-glycan overlap

2G12 is the only known bNAb that targets only glycans^105–107^. Figure 12). Amongst the six clades, clade A and G glycans have minimum occupancy in their bound volume. There are a few glycans in the V1 and V4 loops of clades B and 01 AE. The remaining clades have a substantial occupancy in the bound volume of 2G12, which is not known to be required for 2G12 binding. This suggests that clades A and G might poorly bind to 2G12.

### Variation of the effective number of overlapping atoms

Table S5 and Table S6 in the SI gives the effective number of overlapping glycan atoms, 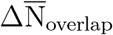 (see methods for definition), for various bNAbs and clades. For a given bNAb, the maximum and minimum values of 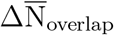 are colored red and blue, respectively. Although 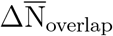 may not directly correlate with the binding efficiency of bNAbs as much as other factors, such as amino acid sequences and loop lengths also influence binding, it gives information about how glycan can interfere with bNAb binding. Table S5 shows that overall, Clade A and 01 AE have maximum effective overlapping glycan atoms for most CD4bs bNAbs, whereas clade B has the least. Interestingly, all V1/V2 apex targeting bNAbs have most favorable interactions with clade B and least favorable interactions with clade 01 AE as shown in Table S6. Overall, many V3-glycan targeting bNAbs have the most favorable interactions with clade A and the least favorable interactions with clade 01 AE and 07 BC.

### Glycan shielding of antibody epitope

An unusually long CDR3 loop allows many bNAbs to penetrate the glycan shield and contact the protein epitopes^108^. We have analyzed whether the epitopes are shielded differently for different clades. Figure 13 displays the avgΔSASA for epitope sites of bNAbs (A) VRC01, (B) 3BNC117, (C) PGT121, and (D) 10-1074 for various clades. The sites affected by glycan shielding are blue with increasing intensity if the shielding is more. Sites absent in the Env structure are shown in black.

**Figure 13.**
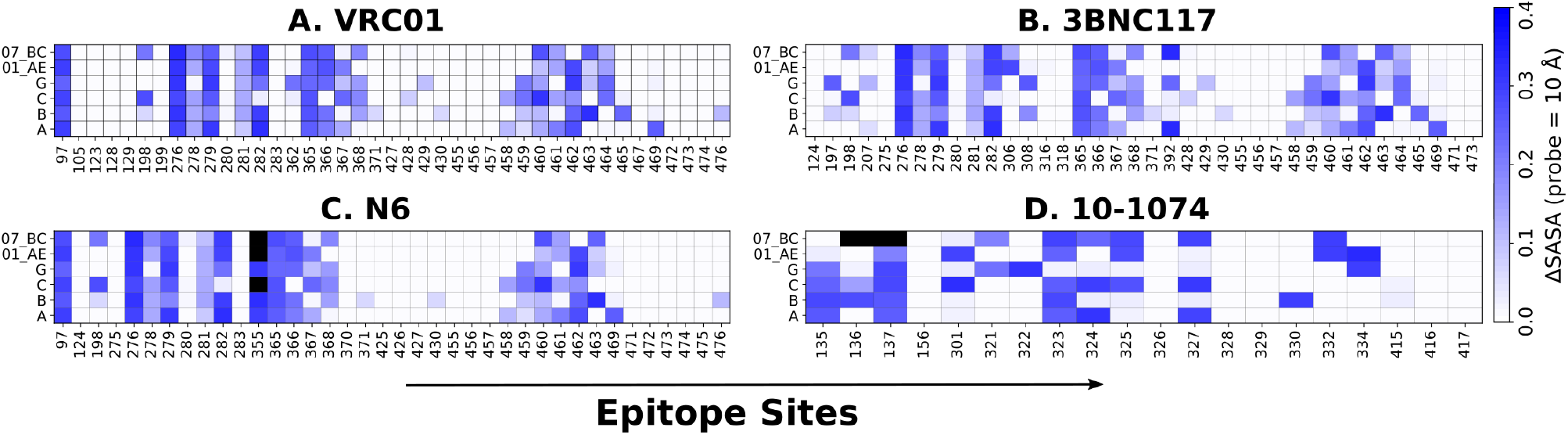
Epitope site (X-axis) ΔSASA for bNAbs (A) VRC01, (B) 3BNC117, (C) N6, (D) 10-1074 for various clades (Y-axis). All ΔSASA are computed with a probe size of 10.0 Å. If an AA site is missing in the crystal structure, it is marked black.

While most sites (in white) are not accessible to the 10.0 Å probe even in the bare protein, glycosylation does affect the epitope sites in a clade-specific manner. For example, epi-tope sites 198, 278,282, 366-368, 458-463 which are common for CD4bs class bNAbs VRC01, 3BNC117, and N6 are differently shielded by glycans. The same is true for epitope sites 136, 324, 334 of V3-glycan class bNAb 10-1074.

To get a quantitative measure of the effect of epitope shielding, we computed the total epitope shielding, 𝒮, for various bNAbs and clades, as shown in Table S7.

For a given bNAb, the maximum and minimum values of 𝒮 are shown in red and blue, respectively. We did not consider shielding due to glycan at N332 for clade A, as it is absent in native clade A variants. We observed that most CD4bs class bNAbs, the total epitope shielding, 𝒮, is minimum for clade 01 AE, suggesting binding of CD4bs class bNAbs is least hindered in clade 01 AE.

All V1/V2 apex class bNAbs considered in this study have the largest 𝒮 for clade 01 AE, whereas clade A has the smallest 𝒮 for most. Epitopes of V1/V2 apex bNAbs are, thus, most accessible in clade A. On the contrary, no pattern is observed for V3-glycan targeting bNAbs. CD4bs bNAbs’s epitopes are shielded most in Clade C and least in Clade 01 AE. Eptipoes of V1/V2 apex bNAbs are shielded the most in clade 01 AE, followed by clade A, 07 BC, B and C in descending order. Epitope shielding of V3glycan bNAbs does not follow a specific pattern but is shielded most in either clade 01 AE or B.

## Conclusion

We find that while glycans cover most of the exposed surface area in all clades, the amount of accessible surface varies, with clade B having the minimum and clade 07 BC the maximum antibody accessible surface area. The number of glycan conformers per glycosylation site also varies with clades, even for conserved sites. We observed that bNAbs interact with more glycans than previously reported in experimental and computational studies ^24,55^. Further, we observe that all the 50 bNAbs have glycans at least PNGS making, on average, more than 10 atoms contacts. Within a clade, almost all bNAbs belonging to a particular epitope class have the same glycans present in their bound volume with a few differences due to the variation in the bNAb’s direction of approach for binding. V2-apex bNAb PGT145 have contacts with glycans from all three protomers of the trimer. Moreover, we find that clade B has the minimum effective overlapping glycan atoms for most CD4bs bNAbs, whereas clades A and 01 AE have the most. All V1/V2 apex targeting bNAbs have the most favorable interactions with clade B and the least favorable interactions with clade 01 AE. In general, epitopes of V1/V2 apex bNAbs are most accessible in clade A, whereas CD4bs bNAbs’ epitopes are shielded most in clade C and least in clade 01 AE. These insights into clade-specific glycan shielding and Env-bNAb interactions at the atomic level will be valuable for improving bNAb-based therapies and informing new vaccine design strategies against HIV-1.

## Limitations of the study

Our focus has been to understand the clade-specific effects of glycans on bNAbs. We did not consider the impact of the type of antibody-antigen interactions and variation in AA residues. Strains within a clade can have variations in the number and positions of PNGS sites. Hence, our analysis is strictly valid for the specific strain of each clade considered here. Molecular dynamics simulations, used to incorporate protein dynamics, were carried out for non-glycosylated HIV-1 Env trimers. The presence of glycans can affect the conformational dynamics of the flexible loop regions. Thus, our analysis might not be accurate if glycans substantially impact the flexible loop conformers. Glycoshield does not take into account glycan-glycan interactions. Glycan-glycan interaction can alter the topology of the glycan cloud; ^24,53^ however, such effects are generally small, and we assume that these would not affect our analysis. In addition, our analysis was performed using Env trimers glycosylated with Man9 glycans. Native-like HIV-1 Envs possess a mixture of high-mannose, hybrid, and complex-type glycans, which can vary across clades. Chakraborty *et. al*.^53^ reported only subtle variations in glycan structures with native-like vs Man9 glycosylation of HIV-1 Env of BG505 (clade A) strain. Our Man9 vs native-like glycosylation of clade A structure for various properties shows only a minor variation, supporting the findings of Chakraborty *et. al*.^53^ and further strengthens the validity of our results. There might be strain-specific differences for glycan effects on bNAb binding^84,92^. In our study, we assumed knowledge of the importance (or not) of individual glycans for bNAb recognition for all clades. However, known experimental results are for a specific strain or clade. In calculating effective epitope shielding, we assumed that epitope sites are identical for all clades. As bNAbs mainly target conserved sites on the Env trimer, our assumption will hold for most epitope sites. Finally, while computing 𝒮, we assumed that all epitope sites are equally important, which is typically not the case.

## Supporting information

The additional data in the Supporting Information file support the results presented in the manuscript.

## Author Information

## Acknowledgments

We thank DST, India for funding under joint Indo-Taiwan program in science and technology (project #2023/IN-TW/02)

## Author Contributions

Mrinal Aranadhara: Conceptualization (equal); Methodology (lead); Formal analysis (lead); Data curation (lead); Visualization (lead); Writing – original draft (lead); Writing – review & editing (equal). Yogendra Kumar: Data curation (supporting); Visualization (supporting); Writing – original draft (MD Simulations). Narendra Dixit: Conceptualization (lead); Supervision (lead); Writing – review & editing (lead). Prabal Maiti: Conceptualization (lead); Supervision (lead); Funding acquisition(lead); Resources (lead), Writing – review & editing (lead).

## Data and Software Availability

This repository contains all the files required to perform the analysis. The information collected here is organized to facilitate the reproduction of the results. The aim of this repository is to serve as a comprehensive and accessible resource for individuals interested in replicating or building upon this work for future research. https://doi.org/10.5281/zenodo.16946212.

## References

(1) West, A.; Scharf, L.; Scheid, J.; Klein, F.; Bjorkman, P.; Nussenzweig, M. Structural Insights on the Role of Antibodies in HIV-1 Vaccine and Therapy. Cell 2014, 156, 633–648.

(2) Korber, B.; Hraber, P.; Wagh, K.; Hahn, B. H. Polyvalent vaccine approaches to combat ¡scp¿HIV¡/scp¿-1 diversity. Immunol. Rev. 2017, 275, 230–244.

(3) Ward, A. B.; Wilson, I. A. The HIV-1 envelope glycoprotein structure: nailing down a moving target. Immunol Rev 2017, 275, 21–32.

(4) Behrens, A.-J. et al. Composition and Antigenic Effects of Individual Glycan Sites of a Trimeric HIV-1 Envelope Glycoprotein. Cell Reports 2016, 14, 2695–2706.

(5) Cao, L. et al. Global site-specific N-glycosylation analysis of HIV envelope glycoprotein. Nat Commun 2017, 8 .

(6) Cao, L. et al. Differential processing of HIV envelope glycans on the virus and soluble recombinant trimer. Nat Commun 2018, 9 .

(7) Watanabe, Y.; Allen, J. D.; Wrapp, D.; McLellan, J. S.; Crispin, M. Site-specific glycan analysis of the SARS-CoV-2 spike. Science 2020, 369, 330–333.

(8) Lasky, L. A.; Groopman, J. E.; Fennie, C. W.; Benz, P. M.; Capon, D. J.; Dowbenko, D. J.; Nakamura, G. R.; Nunes, W. M.; Renz, M. E.; Berman, P. W. Neutralization of the AIDS Retrovirus by Antibodies to a Recombinant Envelope Glycoprotein. Science 1986, 233, 209–212.

(9) Depetris, R. S. et al. Partial Enzymatic Deglycosylation Preserves the Structure of Cleaved Recombinant HIV-1 Envelope Glycoprotein Trimers. J. Biol. Chem. 2012, 287, 24239–24254.

(10) Wagh, K.; Hahn, B. H.; Korber, B. Hitting the sweet spot: exploiting HIV-1 glycan shield for induction of broadly neutralizing antibodies. Current Opinion in HIV and AIDS 2020, 15, 267–274.

(11) Deimel, L. P.; Xue, X.; Sattentau, Q. J. Glycans in HIV-1 vaccine design – engaging the shield. Trends Microbiol 2022, 30, 866–881.

(12) Quinones-Kochs, M. I.; Buonocore, L.; Rose, J. K. Role of N-Linked Glycans in a Human Immunodeficiency Virus Envelope Glycoprotein: Effects on Protein Function and the Neutralizing Antibody Response. Journal of Virology 2002, 76, 4199–4211.

(13) Huang, J. et al. Identification of a CD4-Binding-Site Antibody to HIV that Evolved Near-Pan Neutralization Breadth. Immunity 2016, 45, 1108–1121.

(14) Berndsen, Z. T.; Chakraborty, S.; Wang, X.; Cottrell, C. A.; Torres, J. L.; Diedrich, J. K.; López, C. A.; Yates, J. R.; van Gils, M. J.; Paulson, J. C.; Gnanakaran, S.; Ward, A. B. Visualization of the HIV-1 Env glycan shield across scales. Proceedings of the National Academy of Sciences 2020, 117, 28014–28025.

(15) Newby, M. L.; Allen, J. D.; Crispin, M. Influence of glycosylation on the immunogenicity and antigenicity of viral immunogens. Biotechnol Adv 2024, 70, 108283.

(16) Mouquet, H. Antibody B cell responses in HIV-1 infection. Trends in Immunology 2014, 35, 549–561.

(17) Landais, E.; Moore, P. L. Development of broadly neutralizing antibodies in HIV-1 infected elite neutralizers. Retrovirology 2018, 15 .

(18) Wang, Z. et al. A broadly neutralizing macaque monoclonal antibody against the HIV-1 V3-Glycan patch. eLife 2020, 9 .

(19) Haynes, B. F.; Wiehe, K.; Borrow, P.; Saunders, K. O.; Korber, B.; Wagh, K.; McMichael, A. J.; Kelsoe, G.; Hahn, B. H.; Alt, F.; Shaw, G. M. Strategies for HIV-1 vaccines that induce broadly neutralizing antibodies. Nat. Rev. Immunol. 2022, 23, 142–158.

(20) Santerre, M.; Wang, Y.; Arjona, S.; Allen, C.; Sawaya, B. E. Differential Contribution of HIV-1 Subtypes B and C to Neurological Disorders: Mechanisms and Possible Treatments. AIDS Rev 2019, 21 .

(21) Hemelaar, J. et al. Global and regional molecular epidemiology of HIV-1, 1990–2015: a systematic review, global survey, and trend analysis. The Lancet Infectious Diseases 2019, 19, 143–155.

(22) Bobkov, A. F.; Kazennova, E. V.; Selimova, L. M.; Khanina, T. A.; Ryabov, G. S.; Bobkova, M. R.; Sukhanova, A. L.; Kravchenko, A. V.; Ladnaya, N. N.; Weber, J. N.; Pokrovsky, V. V. Temporal trends in the HIV-1 epidemic in Russia: Predominance of subtype A. J. Med. Virol. 2004, 74, 191–196.

(23) Niu, J.; Wang, Q.; Zhao, W.; Meng, B.; Xu, Y.; Zhang, X.; Feng, Y.; Qi, Q.; Hao, Y.; Zhang, X.; Liu, Y.; Xiang, J.; Shao, Y.; Yang, B. Structures and immune recognition of Env trimers from two Asia prevalent HIV-1 CRFs. Nat Commun 2023, 14 .

(24) Stewart-Jones, G. et al. Trimeric HIV-1-Env Structures Define Glycan Shields from Clades A, B, and G. Cell 2016, 165, 813–826.

(25) Zhou, T. et al. Quantification of the Impact of the HIV-1-Glycan Shield on Antibody Elicitation. Cell Reports 2017, 19, 719–732.

(26) Kwong, P. D.; Mascola, J. R.; Nabel, G. J. Broadly neutralizing antibodies and the search for an HIV-1 vaccine: the end of the beginning. Nat. Rev. Immunol. 2013, 13, 693–701.

(27) Caskey, M.; Klein, F.; Nussenzweig, M. C. Broadly neutralizing anti-HIV-1 monoclonal antibodies in the clinic. Nat. Med. 2019, 25, 547–553.

(28) Walker, L. M.; Burton, D. R. Passive immunotherapy of viral infections: “super-antibodies” enter the fray. Nat. Rev. Immunol. 2018, 18, 297–308.

(29) Nandy, B.; Bindu, D. H.; Dixit, N. M.; Maiti, P. K. Simulations reveal that the HIV-1 gp120-CD4 complex dissociates via complex pathways and is a potential target of the polyamidoamine (PAMAM) dendrimer. The Journal

(30) Nandy, B.; Saurabh, S.; Sahoo, A. K.; Dixit, N. M.; Maiti, P. K. The SPL7013 dendrimer destabilizes the HIV-1 gp120–CD4 complex. Nanoscale 2015, 7, 18628–18641.

(31) Rajendra, D.; Maroli, N.; Dixit, N. M.; Maiti, P. K. Molecular dynamics simulations show how antibodies may rescue HIV-1 mutants incapable of infecting host cells. Journal of Biomolecular Structure and Dynamics 2025, 43, 2982–2992.

(32) Garg, A. K.; Desikan, R.; Dixit, N. M. Preferential presentation of high-affinity immune complexes in germinal centers can explain how passive immunization improves the humoral response. Cell reports 2019, 29, 3946–3957.

(33) Desikan, R.; Raja, R.; Dixit, N. M. Early exposure to broadly neutralizing antibodies may trigger a dynamical switch from progressive disease to lasting control of SHIV infection. PLoS computational biology 2020, 16, e1008064.

(34) Saha, A.; Dixit, N. M. Pre-existing resistance in the latent reservoir can compromise VRC01 therapy during chronic HIV-1 infection. PLOS Computational Biology 2020, 16, e1008434.

(35) LaMont, C.; Otwinowski, J.; Vanshylla, K.; Gruell, H.; Klein, F.; Nourmohammad, A. Design of an optimal combination therapy with broadly neutralizing antibodies to suppress HIV-1. Elife 2022, 11, e76004.

(36) Gorai, B.; Sahoo, A. K.; Srivastava, A.; Dixit, N. M.; Maiti, P. K. Concerted interactions between multiple gp41 trimers and the target cell lipidome may be required for HIV-1 entry. Journal of Chemical Information and Modeling 2020, 61, 444–454.

(37) Gorai, B.; Das, S.; Maiti, P. K. Prediction and validation of HIV-1 gp41 ecto-transmembrane domain post-fusion trimeric structure using molecular modeling. Journal of Biomolecular Structure and Dynamics 2020, 38, 2592–2603.

(38) Crispin, M.; Doores, K. J. Targeting host-derived glycans on enveloped viruses for antibody-based vaccine design. Current Opinion in Virology 2015, 11, 63–69.

(39) Klasse, P.; Ozorowski, G.; Sanders, R. W.; Moore, J. P. Env Exceptionalism: Why Are HIV-1 Env Glycoproteins Atypical Immunogens? Cell Host & Microbe 2020, 27, 507–518.

(40) Wang, S.; Mata-Fink, J.; Kriegsman, B.; Hanson, M.; Irvine, D. J.; Eisen, H. N.; Burton, D. R.; Wittrup, K. D.; Kardar, M.; Chakraborty, A. K. Manipulating the selection forces during affinity maturation to generate cross-reactive HIV antibodies. Cell 2015, 160, 785–797.

(41) Vemparala, B.; Chowdhury, S.; Guedj, J.; Dixit, N. M. Modelling HIV-1 control and remission. NPJ systems biology and applications 2024, 10, 84.

(42) Vemparala, B.; Guedj, J.; Dixit, N. M. Advances in the mathematical modeling of posttreatment control of HIV-1. Current Opinion in HIV and AIDS 2025, 20, 92–98.

(43) Varki, A.; Cummings, R. D.; Esko, J. D.; Stanley, P.; Hart, G. W.; Aebi, M.; Darvill, A. G.; Kinoshita, T.; Packer, N. H.; Prestegard, J. H.; others Essentials of Glycobiology; Cold spring harbor laboratory press, 2015.

(44) Go, E. P.; Chang, Q.; Liao, H.-X.; Sutherland, L. L.; Alam, S. M.; Haynes, B. F.; Desaire, H. Glycosylation Site-Specific Analysis of Clade C HIV-1 Envelope Proteins. J. Proteome Res. 2009, 8, 4231–4242.

(45) Go, E. P.; Hewawasam, G.; Liao, H.-X.; Chen, H.; Ping, L.-H.; Anderson, J. A.; Hua, D. C.; Haynes, B. F.; Desaire, H. Characterization of Glycosylation Profiles of HIV-1 Transmitted/Founder Envelopes by Mass Spectrometry. J. Virol. 2011, 85, 8270–8284.

(46) Go, E. P.; Liao, H.-X.; Alam, S. M.; Hua, D.; Haynes, B. F.; Desaire, H. Characterization of Host-Cell Line Specific Glycosylation Profiles of Early Transmitted/Founder HIV-1 gp120 Envelope Proteins. J. Proteome Res. 2013, 12, 1223–1234.

(47) Go, E. P.; Herschhorn, A.; Gu, C.; Castillo-Menendez, L.; Zhang, S.; Mao, Y.; Chen, H.; Ding, H.; Wakefield, J. K.; Hua, D.; Liao, H.-X.; Kappes, J. C.; Sodroski, J.; Desaire, H. Comparative Analysis of the Glycosylation Profiles of Membrane-Anchored HIV-1 Envelope Glycoprotein Trimers and Soluble gp140. J. Virol. 2015, 89, 8245–8257.

(48) Mouquet, H.; Scharf, L.; Euler, Z.; Liu, Y.; Eden, C.; Scheid, J. F.; Halper-Stromberg, A.; Gnanapragasam, P. N. P.; Spencer, D. I. R.; Seaman, M. S.; Schuitemaker, H.; Feizi, T.; Nussenzweig, M. C.; Bjorkman, P. J. Complex-type N -glycan recognition by potent broadly neutralizing HIV antibodies. PNAS 2012, 109 .

(49) Kong, L. et al. Supersite of immune vulnerability on the glycosylated face of HIV-1 envelope glycoprotein gp120. Nature Structural & Molecular Biology 2013, 20, 796–803.

(50) Falkowska, E. et al. Broadly Neutralizing HIV Antibodies Define a Glycan-Dependent Epitope on the Prefusion Conformation of gp41 on Cleaved Envelope Trimers. Immunity 2014, 40, 657–668.

(51) Gristick, H. B.; von Boehmer, L.; West Jr, A. P.; Schamber, M.; Gazumyan, A.; Golijanin, J.; Seaman, M. S.; Fätkenheuer, G.; Klein, F.; Nussenzweig, M. C.; Bjorkman, P. J. Natively glycosylated HIV-1 Env structure reveals new mode for antibody recognition of the CD4-binding site. Nature Structural &; Molecular Biology 2016, 23, 906–915.

(52) Barnes, C. O.; Gristick, H. B.; Freund, N. T.; Escolano, A.; Lyubimov, A. Y.; Hartweger, H.; West, A. P.; Cohen, A. E.; Nussenzweig, M. C.; Bjorkman, P. J. Structural characterization of a highly-potent V3-glycan broadly neutralizing antibody bound to natively-glycosylated HIV-1 envelope. Nat Commun 2018, 9 .

(53) Chakraborty, S.; Berndsen, Z. T.; Hengartner, N. W.; Korber, B. T.; Ward, A. B.; Gnanakaran, S. Quantification of the Resilience and Vulnerability of HIV-1 Native Glycan Shield at Atomistic Detail. iScience 2020, 23, 101836.

(54) Lemmin, T.; Soto, C.; Stuckey, J.; Kwong, P. D. Microsecond Dynamics and Network Analysis of the HIV-1 SOSIP Env Trimer Reveal Collective Behavior and Conserved Microdomains of the Glycan Shield. Structure 2017, 25, 1631–1639.e2.

(55) Yang, M.; Huang, J.; Simon, R.; Wang, L.-X.; MacKerell, A. D. Conformational Heterogeneity of the HIV Envelope Glycan Shield. Scientific Reports 2017, 7 .

(56) Bennett, A. L.; Edwards, R.; Kosheleva, I.; Saunders, C.; Bililign, Y.; Williams, A.; Bubphamala, P.; Manosouri, K.; Anasti, K.; Saunders, K. O.; Alam, S. M.; Haynes, B. F.; Acharya, P.; Henderson, R. Microsecond dynamics control the HIV-1 Envelope conformation. Sci. Adv. 2024, 10 .

(57) Tsai, Y.-X. et al. Rapid simulation of glycoprotein structures by grafting and steric exclusion of glycan conformer libraries. Cell 2024, 187, 1296–1311.e26.

(58) Wagh, K. et al. Completeness of HIV-1 Envelope Glycan Shield at Transmission Determines Neutralization Breadth. Cell Reports 2018, 25, 893–908.e7.

(59) Shet, A.; Nagaraja, P.; Dixit, N. M. Viral decay dynamics and mathematical modeling of treatment response: Evidence of lower in vivo fitness of HIV-1 subtype C. JAIDS Journal of Acquired Immune Deficiency Syndromes 2016, 73, 245–251.

(60) Rusert, P. et al. Determinants of HIV-1 broadly neutralizing antibody induction. Nat Med 2016, 22, 1260–1267.

(61) Berman, H. M. The Protein Data Bank. Nucleic Acids Research 2000, 28, 235–242.

(62) Henderson, R.; Zhou, Y.; Stalls, V.; Wiehe, K.; Saunders, K. O.; Wagh, K.; Anasti, K.; Barr, M.; Parks, R.; Alam, S. M.; Korber, B.; Haynes, B. F.; Bartesaghi, A.; Acharya, P. Structural basis for breadth development in the HIV-1 V3-glycan targeting DH270 antibody clonal lineage. Nat. Commun. 2023, 14 .

(63) Šali, A.; Blundell, T. L. Comparative Protein Modelling by Satisfaction of Spatial Restraints. Journal of Molecular Biology 1993, 234, 779–815.

(64) Lindahl, E.; Abraham, M. J.; Hess, B.; van der Spoel, D. GROMACS 2019.6. Zenodo 2019,

(65) Zhang, M.; Gaschen, B.; Blay, W.; Foley, B.; Haigwood, N.; Kuiken, C.; Korber, B. Tracking global patterns of N-linked glycosylation site variation in highly variable viral glycoproteins: HIV, SIV, and HCV envelopes and influenza hemagglutinin. Glycobiology 2004, 14, 1229–1246.

(66) Van Der Spoel, D.; Lindahl, E.; Hess, B.; Groenhof, G.; Mark, A. E.; Berendsen, H. J. GROMACS: fast, flexible, and free. Journal of computational chemistry 2005, 26, 1701–1718.

(67) Huang, J.; MacKerell Jr, A. D. CHARMM36 all-atom additive protein force field: Validation based on comparison to NMR data. Journal of computational chemistry 2013, 34, 2135–2145.

(68) Mark, P.; Nilsson, L. Structure and dynamics of the TIP3P, SPC, and SPC/E water models at 298 K. The Journal of Physical Chemistry A 2001, 105, 9954–9960.

(69) NOSÉ, S. U. I. A molecular dynamics method for simulations in the canonical ensemble. Molecular physics 2002, 100, 191–198.

(70) Hoover, W. G. Canonical dynamics: Equilibrium phase-space distributions. Physical review A 1985, 31, 1695.

(71) Parrinello, M.; Rahman, A. Polymorphic transitions in single crystals: A new molecular dynamics method. Journal of Applied physics 1981, 52, 7182–7190.

(72) Hess, B.; Bekker, H.; Berendsen, H. J.; Fraaije, J. G. LINCS: A linear constraint solver for molecular simulations. Journal of computational chemistry 1997, 18, 1463–1472.

(73) Darden, T.; York, D.; Pedersen, L.; others Particle mesh Ewald: An N log (N) method for Ewald sums in large systems. Journal of chemical physics 1993, 98, 10089–10089.

(74) Essmann, U.; Perera, L.; Berkowitz, M. L.; Darden, T.; Lee, H.; Pedersen, L. G. A smooth particle mesh Ewald method. The Journal of chemical physics 1995, 103, 8577–8593.

(75) Meng, E. C.; Pettersen, E. F.; Couch, G. S.; Huang, C. C.; Ferrin, T. E. Tools for integrated sequence-structure analysis with UCSF Chimera. BMC Bioinformatics 2006, 7, 339, Matchmaker program implemented in UCSF Chimera and later incorporated into ChimeraX.

(76) Pettersen, E. F.; Goddard, T. D.; Huang, C. C.; Meng, E. C.; Couch, G. S.; Croll, T. I.; Morris, J. H.; Ferrin, T. E. UCSF ChimeraX: Structure visualization for researchers, educators, and developers. Protein Science 2021, 30, 70–82.

(77) Los Alamos National Laboratory Neutralizing Antibody Contacts & Features Database. 2025; https://www.hiv.lanl.gov/content/immunology, xAccessed: July 14, 2024.

(78) Wibmer, C. K.; Moore, P. L.; Morris, L. HIV broadly neutralizing antibody targets. Current Opinion in HIV and AIDS 2015, 10, 135–143.

(79) Sok, D.; Burton, D. R. Recent progress in broadly neutralizing antibodies to HIV. Nat Immunol 2018, 19, 1179–1188.

(80) Pancera, M. et al. Structure and immune recognition of trimeric pre-fusion HIV-1 Env. Nature 2014, 514, 455–461.

(81) Grant, O. C.; Montgomery, D.; Ito, K.; Woods, R. J. Analysis of the SARS-CoV-2 spike protein glycan shield reveals implications for immune recognition. Scientific Reports 2020, 10 .

(82) Sanders, R. W. et al. A Next-Generation Cleaved, Soluble HIV-1 Env Trimer, BG505 SOSIP.664 gp140, Expresses Multiple Epitopes for Broadly Neutralizing but Not Non-Neutralizing Antibodies. PLoS Pathogens 2013, 9, e1003618.

(83) Dingens, A. S.; Arenz, D.; Weight, H.; Overbaugh, J.; Bloom, J. D. An Antigenic Atlas of HIV-1 Escape from Broadly Neutralizing Antibodies Distinguishes Functional and Structural Epitopes. Immunity 2019, 50, 520–532.e3.

(84) Bricault, C. A. et al. HIV-1 Neutralizing Antibody Signatures and Application to Epitope-Targeted Vaccine Design. Cell Host & Microbe 2019, 25, 59–72.e8.

(85) Georgiev, I. S.; Gordon Joyce, M.; Zhou, T.; Kwong, P. D. Elicitation of HIV-1-neutralizing antibodies against the CD4-binding site. Current Opinion in HIV and AIDS 2013, 8, 382–392.

(86) Mahomed, S.; Garrett, N.; Baxter, C.; Abdool Karim, Q.; Abdool Karim, S. S. Clinical Trials of Broadly Neutralizing Monoclonal Antibodies for Human Immunodeficiency Virus Prevention: A Review. The Journal of Infectious Diseases 2020, 223, 370–380.

(87) Taiwo, B.; others VH3810109 (N6LS) Efficacy and Safety in Adults Who Are Virologically Suppressed: The EMBRACE Study. Conference on Retroviruses and Opportunistic Infections (CROI 2025). San Francisco, CA, 2025; Presented March 9–12.

(88) van Schooten, J. et al. Identification of IOMA-class neutralizing antibodies targeting the CD4-binding site on the HIV-1 envelope glycoprotein. Nature Communications 2022, 13 .

(89) Andrabi, R.; Voss, J.; Liang, C.-H.; Briney, B.; McCoy, L.; Wu, C.-Y.; Wong, C.-H.; Poignard, P.; Burton, D. Identification of Common Features in Prototype Broadly Neutralizing Antibodies to HIV Envelope V2 Apex to Facilitate Vaccine Design. Immunity 2015, 43, 959–973.

(90) Doria-Rose, N. A. et al. Developmental pathway for potent V1V2-directed HIV-neutralizing antibodies. Nature 2014, 509, 55–62.

(91) McLellan, J. S. et al. Structure of HIV-1 gp120 V1/V2 domain with broadly neutralizing antibody PG9. Nature 2011, 480, 336–343.

(92) Doria-Rose, N. A. et al. New Member of the V1V2-Directed CAP256-VRC26 Lineage That Shows Increased Breadth and Exceptional Potency. J Virol 2016, 90, 76–91.

(93) Julien, J.-P.; Lee, J. H.; Cupo, A.; Murin, C. D.; Derking, R.; Hoffenberg, S.; Caulfield, M. J.; King, C. R.; Marozsan, A. J.; Klasse, P. J.; Sanders, R. W.; Moore, J. P.; Wilson, I. A.; Ward, A. B. Asymmetric recognition of the HIV-1 trimer by broadly neutralizing antibody PG9. Proceedings of the National Academy of Sciences 2013, 110, 4351–4356.

(94) Freund, N. T. et al. Coexistence of potent HIV-1 broadly neutralizing antibodies and antibody-sensitive viruses in a viremic controller. Science Translational Medicine 2017, 9, eaal2144.

(95) Scharf, L.; Wang, H.; Gao, H.; Chen, S.; McDowall, A.; Bjorkman, P. Broadly Neutralizing Antibody 8ANC195 Recognizes Closed and Open States of HIV-1 Env. Cell 2015, 162, 1379–1390.

(96) Kong, L.; Torrents de la Peña, A.; Deller, M. C.; Garces, F.; Sliepen, K.; Hua, Y.; Stanfield, R. L.; Sanders, R. W.; Wilson, I. A. Complete epitopes for vaccine design derived from a crystal structure of the broadly neutralizing antibodies PGT128 and 8ANC195 in complex with an HIV-1 Env trimer. Acta Crystallographica Section D Biological Crystallography 2015, 71, 2099–2108.

(97) Kong, R. et al. Fusion peptide of HIV-1 as a site of vulnerability to neutralizing antibody. Science 2016, 352, 828–833.

(98) Cottrell, C. A. et al. Structural basis of glycan276-dependent recognition by HIV-1 broadly neutralizing antibodies. Cell Reports 2021, 37, 109922.

(99) Scharf, L.; Scheid, J.; Lee, J.; West, A.; Chen, C.; Gao, H.; Gnanapragasam, P.; Mares, R.; Seaman, M.; Ward, A.; Nussenzweig, M.; Bjorkman, P. Antibody 8ANC195 Reveals a Site of Broad Vulnerability on the HIV-1 Envelope Spike. Cell Reports 2014, 7, 785–795.

(100) Huang, J. et al. Broad and potent HIV-1 neutralization by a human antibody that binds the gp41–gp120 interface. Nature 2014, 515, 138–142.

(101) Do Kwon, Y. et al. Crystal structure, conformational fixation and entryrelated interactions of mature ligandfree HIV-1 Env. Nature Structural & Molecular Biology 2015, 22, 522–531.

(102) Blattner, C. et al. Structural Delineation of a Quaternary, Cleavage-Dependent Epitope at the gp41-gp120 Interface on Intact HIV-1 Env Trimers. Immunity 2014, 40, 669–680.

(103) Dingens, A. S.; Haddox, H. K.; Overbaugh, J.; Bloom, J. D. Comprehensive Mapping of HIV-1 Escape from a Broadly Neutralizing Antibody. Cell Host & Microbe 2017, 21, 777–787.e4.

(104) Lee, J. H.; Ozorowski, G.; Ward, A. B. Cryo-EM structure of a native, fully glycosylated, cleaved HIV-1 envelope trimer. Science 2016, 351, 1043–1048.

(105) Pantophlet, R.; Ollmann Saphire, E.; Poignard, P.; Parren, P. W. H. I.; Wilson, I. A.; Burton, D. R. Fine Mapping of the Interaction of Neutralizing and Nonneutralizing Monoclonal Antibodies with the CD4 Binding Site of Human Immunodeficiency Virus Type 1 gp120. J Virol 2003, 77, 642–658.

(106) Alam, S. M. et al. Mimicry of an HIV broadly neutralizing antibody epitope with a synthetic glycopeptide. Sci Transl Med 2017, 9 .

(107) Chuang, G.-Y. et al. Structural Survey of Broadly Neutralizing Antibodies Targeting the HIV-1 Env Trimer Delineates Epitope Categories and Characteristics of Recognition. Structure 2019, 27, 196–206.e6.

(108) Doores, K. J. The ¡scp¿HIV¡/scp¿ glycan shield as a target for broadly neutralizing antibodies. The FEBS Journal 2015, 282, 4679–4691.

