## Supplementary material for "Clade-specific influences of glycans on the interactions between HIV-1 envelope and broadly neutralizing antibodies": The additional data in the Supporting Information file support the results presented in the manuscript.

### List of Tables

S1 Schematic structure and composition of various glycoforms. The last column lists the corresponding glycosylation sites (for clade A) used to build native-like glycosylation patterns. Information about native-like glycans are collected from ref.[1] Please note that we used native-like glycan pattern only for clade A to compare our key results in the main mainscript, where Man9 glycoform was used for generating all the glycosylated models of HIV-1 trimer. Fucosylated two-antennae glycan for gp41 and gp120 glycan sites (denoted FA2 in ref[1]) are called A2F and A1F here. Note that due to the unavailability of fucosylated three-antennae complexes (FA3 in ref[1]) in the glycan library of GlycoSHIELD, we replaced glycans sites with FA3 by A2F in our calculations. Green Circle: $\alpha$- and $\beta$- Mannose (Man), Blue Square: N-Acetyl Glucosamine (GlcNAc), Yellow Circle: $\beta$-Galactose (Gal), Red Rhombus: $\alpha$-Fucose (Fuc), and Purple Rhombus: Sialic Acid (SA). . . . . . . . . . . . . . . . . . . . . . . . . . S9

S2 PDB ID of bNAbs of various epitope class used for overlap analysis. We have a total of 30 CD4bs, seven V2-apex, nine V3-glycan class,two gp120/gp41 Interface, and one each of fusion peptide and glycan class bNAbs. . . . . . . S10

S3 Number of glycan conformers ($N_{Acc}$) accepted by GS out of 3000 conformers for various clades (averaged over the 12 fully-glycosylated trimer geometries). Site N339 in clade 01_AE has lowest accepted conformers with $N_{Acc}$ = 196, whereas site N356 in clade 07_BC have the highest, with $N_{Acc}$ = 2914.. Value of $N_{Acc}$ also qualitatively measures glycan conformational dynamics, with larger values corresponding to more volume for glycans to move. Dash denotes missing sites. . . . . . . . . . . . . . . . . . . . . . . . . . . . . . . S11

S4 Number of glycans with at least 10/50 atoms inside the bound volume of various bNAbs. Numbers inside the simple brackets represent the number of protomers the contact glycans belong to. . . . . . . . . . . . . . . . . . . . . S14

S5 Effective number of overlaping glycan atoms, $\Delta\overline{\mathrm{N}}_{\mathrm{overlap}}$, for CD4bs bNAbs. For each bNAb, the minimum and maximum values are colored blue and red, respectively. Clade B has the most favorable interaction with glycans for most of the bNAbs in this class, indicating the binding of CD4bs will be least hindered by glycans. . . . . . . . . . . . . . . . . . . . . . . . . . . . . S16

S6 Effective number of overlaping glycan atoms, $\Delta\overline{\mathrm{N}}_{\mathrm{overlap}}$, for V1/V2 apex (V2), V3-glycan (V3), gp120/gp41 interface (Interface), Fusion Peptide (FP) bNAbs. For each bNAb, the minimum and maximum values are colored blue and red, respectively. Clade B has the most favourable interactions for all of V1/V2 apex bNAbs. Depending on the bNAb, either clade 01_AE or 07_BC has the least favorable glycan interaction for V3-glycan class bNAbs. 2G12, which only target, has many overlapping glycan atoms in clades 01_AE or 07_BC that are not known to be important for its binding or recognition. . . S17

S7 Values of total epitope shielding, $\mathcal{S}$, for various bNAbs. $\mathcal{S}$ gives a measure of how the protein epitopes of bNAbs are shielded by glycans. . . . . . . . . . . S19

S8 Total accessible surface area (Å$^2$) for with Man9 and native-like fully glycosylated HIV-1 Envs for clade A with a probe radii of 1.4, 7.2, and 10.0Å. see Table 1 for site-specific glycoforms used in native-like glycosylation. . . . . . S20

### List of Figures

S1 Structure of clade B (PDB 5FYK) [left: Top View, Right: Side View] highlighting the locations of various Ab epitopes: CD4bs Ab epitope (red), V1/V2 Ab epitope (green), V3 loop Ab epitope (blue), fusion peptide (FP) Ab epitopes (purple), and gp120/gp41 interface Ab epitope (Orange). . . . . . . . . S5

S2 3D structures of various high-mannose (Man5, Man7, Man7, Man9), hybrid (FH), and complex glycans (A1F, A2F). Associated schematic diagrams are shown in Table S1. Green: $\alpha$- and $\beta$- Mannose ( $\alpha$ or $\beta$- Man), Blue: N-Acetyl Glucosamine (GlcNAc), Yellow: $\beta$-Galactose ($\beta$-Gal), Red: $\alpha$-Fucose ($\alpha$-Fuc), and Purple: Sialic Acid (SA). . . . . . . . . . . . . . . . . . . . . . . . . . S6

S3 Gp120 HIV-1 segment (chain G, red) of bNAb VRC01 (chains H, L) is aligned to the corresponding gp120 segments, i.e., chains D, E, and F (shown in light gray) of the modeled structure of BG505 (Clade A, PDBID: 5FYL). . . . . S7

S4 The effect of native-like glycosylation on bNAb-glycan overlap. (a) For CD4bs, V3,FP, Interface and (b) for V2-apex. Comparing with Figures 7, 8, 11, 12, 13 in the main text shows negligible difference between Man9 vs native-like glycans. . . . . . . . . . . . . . . . . . . . . . . . . . . . . . . . . . . . . S8

S5 (A) Structure of Clade B trimer (white) bound to PGT151 (orange) bNAb. Modeled Man9 glycans at key PNGS are also shown. (B) 300 glycan conformers at each PNGS generated using GlycoSHIELD are shown to illustrate PGT151-glycan overlap. PGT151 has significant overlap with glycans at N276, N616, and N637 of one protomer and aslo has contacts with glycans at N262 and N448 of the adjascent protomer. Only a very few atoms of N241 glycans are near the bound PGT151 bNAb. . . . . . . . . . . . . . . . . . . S18

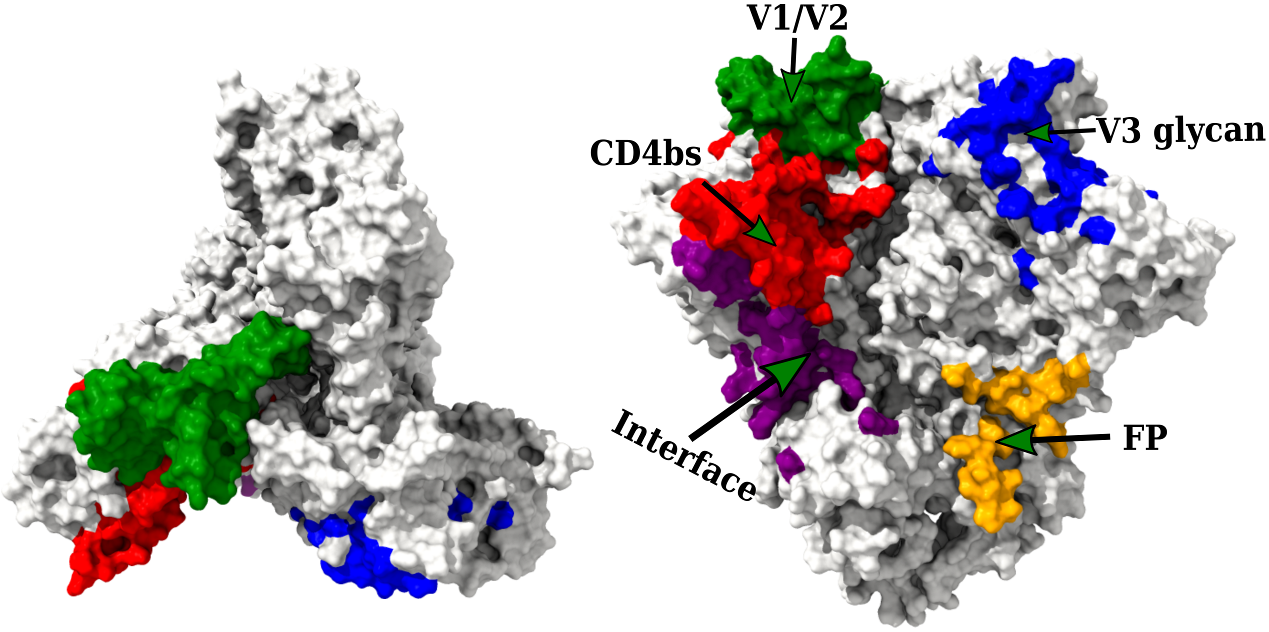


Figure S1: Structure of clade B (PDB 5FYK) [left: Top View, Right: Side View] highlighting the locations of various Ab epitopes: CD4bs Ab epitope (red), V1/V2 Ab epitope (green), V3 loop Ab epitope (blue), fusion peptide (FP) Ab epitopes (purple), and gp120/gp41 interface Ab epitope (Orange).

Figure S2: 3D structures of various high-mannose (Man5, Man7, Man7, Man9), hybrid (FH), and complex glycans (A1F, A2F). Associated schematic diagrams are shown in Table S1. Green: $\alpha$- and $\beta$- Mannose ( $\alpha$ or $\beta$- Man), Blue: N-Acetyl Glucosamine (GlcNAc), Yellow: $\beta$-Galactose ($\beta$-Gal), Red: $\alpha$-Fucose ($\alpha$-Fuc), and Purple: Sialic Acid (SA).

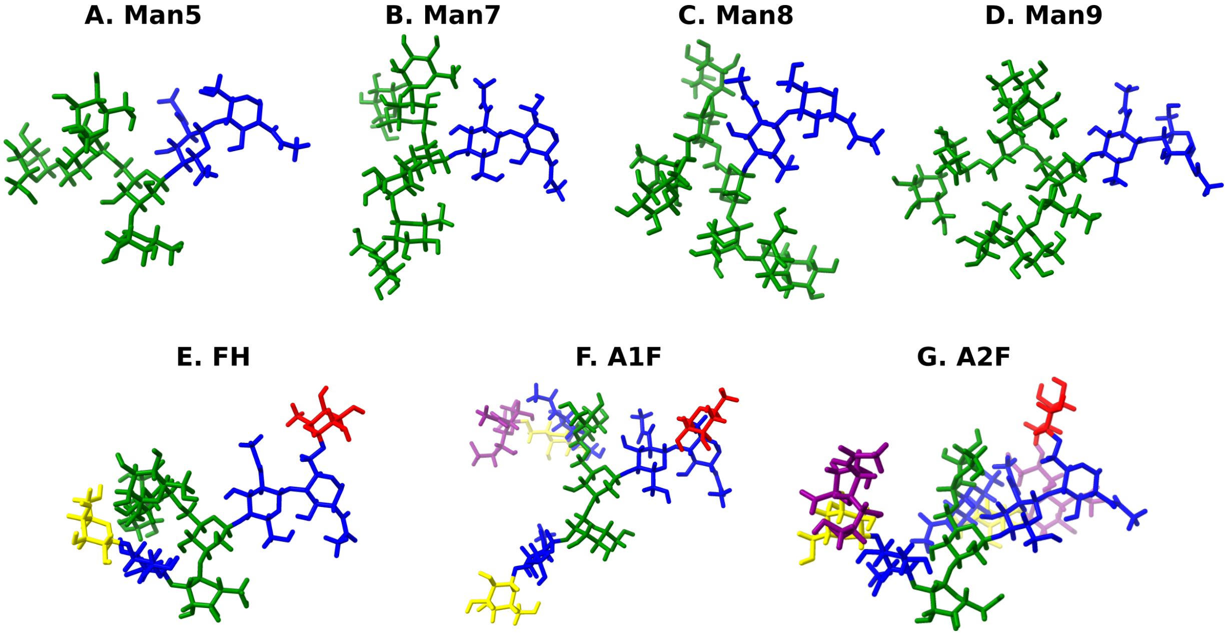

#### Schematic Description of Overlap Analysis

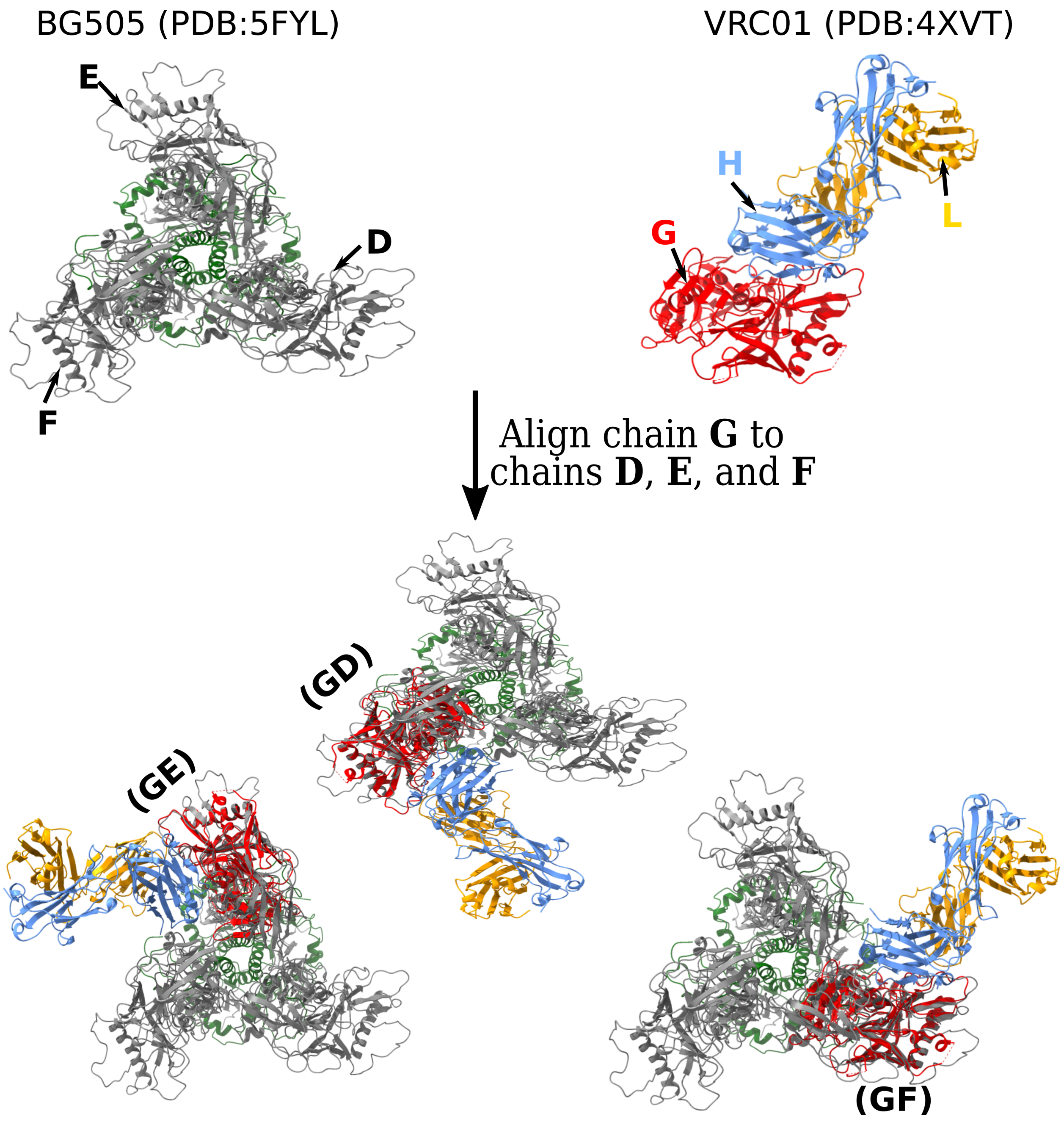


Figure S3: Gp120 HIV-1 segment (chain G, red) of bNAb VRC01 (chains H, L) is aligned to the corresponding gp120 segments, i.e., chains D, E, and F (shown in light gray) of the modeled structure of BG505 (Clade A, PDBID: 5FYL).

#### Glycan-bNAb Overlap Analysis

To analyze the effect of native-like glycosylation, we performed bNAb-glycan overlap analysis for fully-glycosylated clade A trimer. We only find minor differences in the overlap analysis results between Man9 and native-like glycans, indicating that Man9 glycosylation can accurately capture most features of overlap analysis.

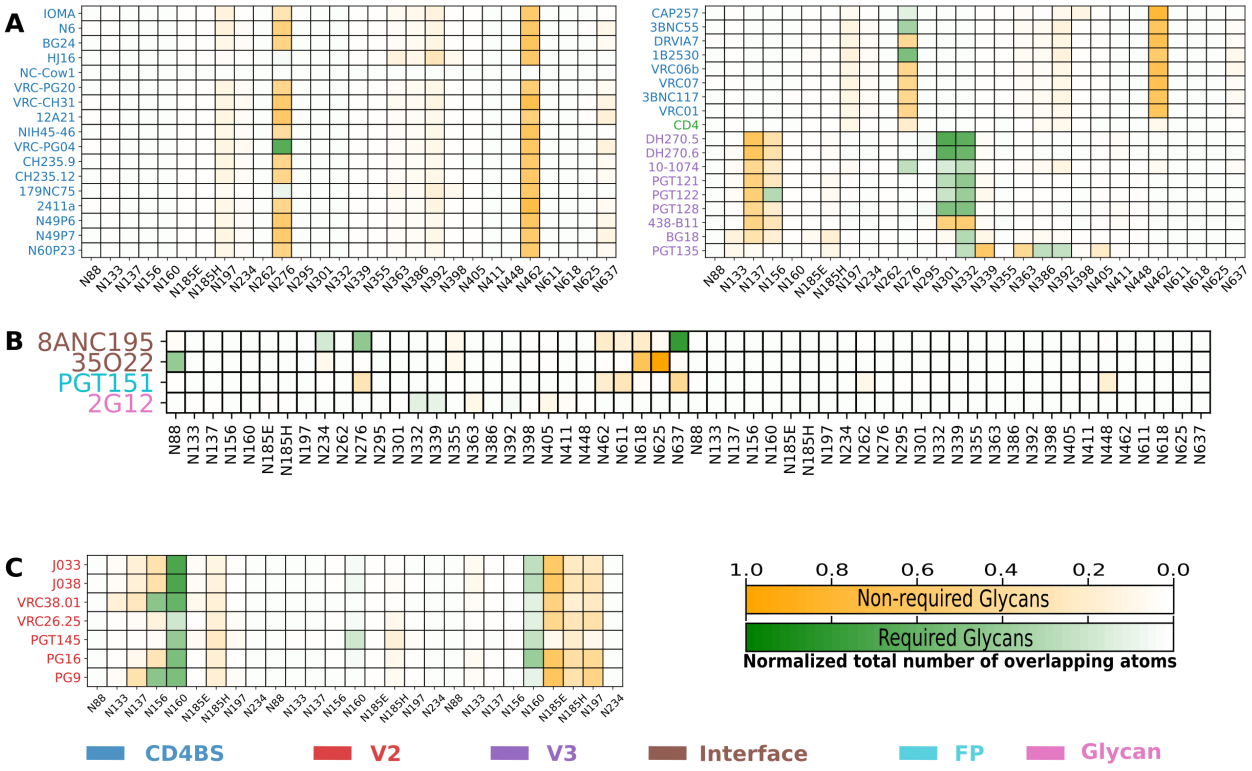


Figure S4: The effect of native-like glycosylation on bNAb-glycan overlap. (a) For CD4bs, V3,FP, Interface and (b) for V2-apex. Comparing with Figures 7, 8, 11, 12, 13 in the main text shows negligible difference between Man9 vs native-like glycans.

Table S1: Schematic structure and composition of various glycoforms. The last column lists the corresponding glycosylation sites (for clade A) used to build native-like glycosylation patterns. Information about native-like glycans are collected from ref.[1] Please note that we used native-like glycan pattern only for clade A to compare our key results in the main mainscript, where Man9 glycoform was used for generating all the glycosylated models of HIV-1 trimer. Fucosylated two-antennae glycan for gp41 and gp120 glycan sites (denoted FA2 in ref[1]) are called A2F and A1F here. Note that due to the unavailability of fucosylated three-antennae complexes (FA3 in ref[1]) in the glycan library of GlycoSHIELD, we replaced glycans sites with FA3 by A2F in our calculations. Green Circle: $\alpha$- and $\beta$- Mannose (Man), Blue Square: N-Acetyl Glucosamine (GlcNAc), Yellow Circle: $\beta$-Galactose (Gal), Red Rhombus: $\alpha$-Fucose (Fuc), and Purple Rhombus: Sialic Acid (SA).

| Name | Glycan Type | Structure | Composition and Glycan Sites |
|---|---|---|---|
| Man5 | High Mannose | 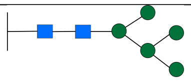 | GlcNAc(2),Man(5)<br>Sites : None |
| Man7 | High Mannose | 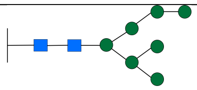 | GlcNAc(2),Man(7)<br>Sites : N276 |
| Man8 | High Mannose | 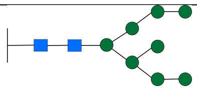 | GlcNAc(2),Man(8)<br>Sites :N160 |
| Man9 | High Mannose | 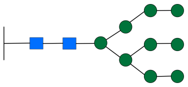 | GlcNAc(2),Man(9)<br>Sites : N133, N156, N234, N262, N295, N301, N332, M339, N363, N386, N392, N411, N448 |
| FH | Hybrid | 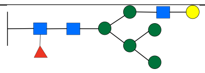 | GlcNAc(2),Man(5)<br>Sites : N355 |
| A2F | Complex | 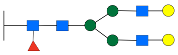 | GlcNAc(4),Man(3),Gal(2),SA(2),Fuc(1)<br>Sites : N611, N618, N625, N637 |
| A1F | Complex | 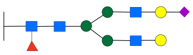 | GlcNAc(4),Man(3),Gal(2),SA(1),Fuc(1)<br>Sites : N88, N137, N185e, N185h, N197, N398, N406, N462 |

Table S2: PDB ID of bNAbs of various epitope class used for overlap analysis. We have a total of 30 CD4bs, seven V2-apex, nine V3-glycan class,two gp120/gp41 Interface, and one each of fusion peptide and glycan class bNAbs.

| bNAb | PDBID | Epitope | bNAb | PDBID | Epitope |
|---|---|---|---|---|---|
| VRC13 | 4YDJ | CD4bs | CD4 | 2QAD | CD4bs |
| BG24 | 7UCG | CD4bs | N6 | 5TE7 | CD4bs |
| VRC-PG04 | 3SE9 | CD4bs | b12 | 2NY7 | CD4bs |
| NIH45-46 | 4JDT | CD4bs | CH103 | 4JAN | CD4bs |
| 3BNC117 | 4JPV | CD4bs | HJ16 | 4YE4 | CD4bs |
| 12A21 | 4JPW | CD4bs | VRC26.25 | 6VTT | V2-apex |
| VRC-CH31 | 4LSP | CD4bs | J038 | 7MXD | V2-apex |
| VRC-PG20 | 4LSU | CD4bs | J033 | 7N28 | V2-apex |
| VRC07 | 4OLZ | CD4bs | PGT145 | 6NIJ | V2-apex |
| 8ANC131 | 4RWY | CD4bs | PG9 | 3U2S | V2-apex |
| VRC02 | 4RX4 | CD4bs | VRC38.01 | 5VGJ | V2-apex |
| VRC06b | 4XNZ | CD4bs | PG16 | 6ULC | V2-apex |
| 1B2530 | 4YFL | CD4bs | PGT135 | 4JM2 | V3-glycan |
| DRVIA7 | 5CD5 | CD4bs | DH270.6 | 6UM6 | V3-glycan |
| CH235.12 | 5F96 | CD4bs | 438-B11 | 6UTK | V3-glycan |
| CH235.9 | 5F9O | CD4bs | DH270.5 | 8SAS | V3-glycan |
| 3BNC55 | 5I9Q | CD4bs | PGT122 | 4TVP | V3-glycan |
| CAP257 | 5T33 | CD4bs | PGT121 | 5CEZ | V3-glycan |
| N60P23 | 5WB9 | CD4bs | 10-1074 | 7UCG | V3-glycan |
| VRC01 | 4XVT | CD4bs | BG18 | 6DFH | V3-glycan |
| N49P7 | 6BCK | CD4bs | PGT128 | 3TYG | V3-glycan |
| IOMA | 5T3Z | CD4bs | 35O22 | 4TVP | gp120/gp41 Interface |
| NC-Cow1 | 6OPA | CD4bs | 8ANC195 | 4P9H | gp120/gp41 Interface |
| N49P6 | 6OZ2 | CD4bs | PGT151 | 6DCQ | gp120/gp41 Interface |
| 2411a | 7JKS | CD4bs | 2G12 | 6OZC | glycan |
| 179NC75 | 7LLK | CD4bs | | | |

Table S3: Number of glycan conformers ($N_{Acc}$) accepted by GS out of 3000 conformers for various clades (averaged over the 12 fully-glycosylated trimer geometries). Site N339 in clade 01_AE has lowest accepted conformers with $N_{Acc}$ = 196, whereas site N356 in clade 07_BC have the highest, with $N_{Acc}$ = 2914.. Value of $N_{Acc}$ also qualitatively measures glycan conformational dynamics, with larger values corresponding to more volume for glycans to move. Dash denotes missing sites.

| PNGS | A | B | C | G | 01_AE | 07_BC |
|---|---|---|---|---|---|---|
| N88 | 2559 | 2442 | 2569 | 2626 | 2755 | 2721 |
| N130 | - | - | - | - | 2123 | - |
| N133 | 1746 | - | 1414 | 1608 | - | - |
| N135 | - | 1139 | - | - | 2056 | - |
| N135B | - | - | - | - | - | 1938 |
| N135F | - | - | - | - | - | 2295 |
| N135I | - | - | - | - | - | 2009 |
| N135O | - | - | - | - | - | 2086 |
| N137 | 2191 | - | - | 779 | - | - |
| N138 | - | 1578 | 2604 | - | - | - |
| N138A | - | - | - | - | 236 | - |
| N138B | - | - | - | 1630 | - | - |
| N138G | - | - | - | - | 1904 | - |
| N138K | - | - | - | - | 2355 | - |
| N141 | - | 1568 | - | - | - | - |
| N142 | - | - | - | 1809 | - | - |
| N149 | - | - | - | - | 1192 | - |
| N156 | 496 | 373 | 1107 | 479 | 381 | 394 |
| N160 | 1721 | 2335 | 2041 | 1726 | 2237 | 2358 |
| N185E | 1534 | - | - | - | - | - |
| N185H | 1415 | - | - | - | - | - |
| N187 | - | - | 1813 | - | 2793 | 2798 |

| PNGS | A | B | C | G | 01_AE | 07_BC |
|---|---|---|---|---|---|---|
| N188 | - | 2656 | - | 2711 | - | - |
| N197 | 514 | - | 630 | 344 | 686 | 486 |
| N234 | 1134 | - | 1694 | 1867 | 1180 | 1263 |
| N241 | - | 2506 | 2529 | 2558 | 2594 | 2562 |
| N262 | 297 | 374 | 370 | 307 | 359 | 348 |
| N276 | 2393 | 2159 | 2364 | 2382 | 2148 | 2302 |
| N289 | - | - | - | - | 201 | 203 |
| N293 | - | - | - | 2611 | - | - |
| N295 | 2173 | 2494 | - | - | - | - |
| N301 | 2462 | 2335 | 2304 | 2472 | 1971 | 2169 |
| N332 | 1512 | 1510 | 1519 | 1192 | - | 1449 |
| N334 | - | - | - | - | 2124 | - |
| N339 | 1017 | 709 | 874 | - | 196 | - |
| N344 | - | - | - | 2331 | - | - |
| N355 | 1761 | 2498 | - | 2497 | - | - |
| N356 | - | - | 2908 | - | 2882 | 2914 |
| N362 | - | 1199 | 1569 | - | - | - |
| N363 | 762 | - | - | - | - | - |
| N386 | 581 | 504 | 595 | 518 | 489 | 705 |
| N392 | 1182 | 1125 | 1101 | 783 | - | 745 |
| N396 | - | - | 2126 | - | - | - |
| N397 | - | 690 | - | - | - | - |
| N398 | 2346 | - | - | - | - | - |
| N402 | - | - | - | - | - | 1691 |
| N403 | - | 1512 | 1219 | - | - | - |
| N404 | - | - | - | - | 1169 | - |
| N405 | 1704 | - | - | 735 | - | - |

| PNGS | A | B | C | G | 01_AE | 07_BC |
|---|---|---|---|---|---|---|
| N406 | - | - | - | - | - | 2206 |
| N408 | - | - | - | 2520 | - | - |
| N409 | - | - | 2469 | - | - | - |
| N411 | 2749 | - | - | - | 2489 | 1939 |
| N413 | - | - | - | 2375 | - | - |
| N442 | - | - | 1130 | 1188 | - | 1034 |
| N444 | - | - | - | - | 1883 | - |
| N448 | 2147 | 2166 | 2375 | - | 2409 | 2312 |
| N462 | 2854 | - | - | 2611 | 2911 | - |
| N463 | - | 2729 | 1721 | - | - | 2685 |
| N464A | - | - | - | 2544 | - | - |
| N611 | 516 | 566 | 1246 | 627 | 1009 | 1203 |
| N616 | - | 2082 | - | 2338 | 2225 | 2231 |
| N618 | 2802 | - | 2787 | - | - | - |
| N625 | 2618 | 2658 | 2745 | 2696 | 2765 | 2763 |
| N637 | 2177 | 1947 | 2493 | 2217 | 2610 | 2661 |

Table S4: Number of glycans with at least 10/50 atoms inside the bound volume of various bNAbs. Numbers inside the simple brackets represent the number of protomers the contact glycans belong to.

| Antibody | A | | B | | C | | G | | 01_AE | | 07_BC | |
|---|---|---|---|---|---|---|---|---|---|---|---|---|
| | >10 | >50 | >10 | >50 | >10 | >50 | >10 | >50 | >10 | >50 | >10 | >50 |
| VRC13 | 12 (2) | 5 (1) | 8 (2) | 3 (2) | 12 (2) | 5 (1) | 11 (2) | 4 (2) | 14 (2) | 6 (2) | 13 (2) | 5 (1) |
| b12 | 12 (2) | 6 (2) | 11 (2) | 3 (2) | 13 (2) | 6 (2) | 14 (2) | 6 (2) | 16 (2) | 10 (2) | 16 (2) | 9 (2) |
| CH103 | 9 (2) | 5 (1) | 8 (2) | 3 (1) | 8 (1) | 5 (1) | 9 (2) | 4 (1) | 8 (2) | 4 (2) | 7 (2) | 5 (1) |
| BG24 | 5 (1) | 2 (1) | 3 (1) | 1 (1) | 5 (1) | 2 (1) | 6 (1) | 1 (1) | 4 (2) | 2 (1) | 5 (1) | 2 (1) |
| HJ16 | 5 (1) | 2 (1) | 5 (1) | 3 (1) | 5 (1) | 3 (1) | 3 (1) | 2 (1) | 3 (1) | 1 (1) | 4 (1) | 2 (1) |
| NC-Cow1 | 8 (2) | 0 (0) | 3 (1) | 0 (0) | 4 (2) | 0 (0) | 5 (2) | 0 (0) | 8 (2) | 0 (0) | 4 (2) | 0 (0) |
| VRC02 | 9 (2) | 2 (1) | 6 (2) | 1 (1) | 9 (2) | 2 (1) | 7 (2) | 1 (1) | 15 (2) | 4 (2) | 11 (2) | 2 (1) |
| 8ANC131 | 9 (2) | 2 (1) | 7 (2) | 1 (1) | 11 (2) | 2 (1) | 7 (2) | 1 (1) | 15 (2) | 4 (2) | 11 (2) | 2 (1) |
| VRC-PG20 | 5 (1) | 2 (1) | 3 (1) | 1 (1) | 5 (1) | 2 (1) | 5 (1) | 1 (1) | 4 (2) | 2 (1) | 5 (1) | 2 (1) |
| VRC-CH31 | 6 (1) | 2 (1) | 5 (1) | 1 (1) | 5 (1) | 2 (1) | 6 (1) | 1 (1) | 7 (2) | 2 (1) | 6 (1) | 2 (1) |
| 12A21 | 6 (2) | 2 (1) | 5 (2) | 1 (1) | 7 (2) | 2 (1) | 6 (2) | 1 (1) | 9 (2) | 2 (1) | 6 (2) | 2 (1) |
| NIH45-46 | 4 (1) | 2 (1) | 4 (1) | 0 (0) | 5 (1) | 2 (1) | 4 (1) | 1 (1) | 4 (2) | 2 (1) | 4 (1) | 2 (1) |
| VRC-PG04 | 5 (1) | 2 (1) | 4 (1) | 1 (1) | 6 (1) | 2 (1) | 5 (1) | 1 (1) | 8 (2) | 2 (1) | 5 (1) | 2 (1) |
| CH235.9 | 4 (1) | 2 (1) | 4 (2) | 1 (1) | 6 (2) | 2 (1) | 5 (2) | 1 (1) | 8 (2) | 2 (1) | 6 (2) | 2 (1) |
| CH235.12 | 4 (1) | 2 (1) | 3 (1) | 1 (1) | 5 (1) | 2 (1) | 4 (1) | 1 (1) | 5 (2) | 2 (1) | 4 (1) | 2 (1) |
| 179NC75 | 5 (1) | 1 (1) | 6 (2) | 1 (1) | 5 (1) | 2 (1) | 3 (2) | 1 (1) | 5 (2) | 1 (1) | 4 (2) | 1 (1) |
| 2411a | 4 (1) | 2 (1) | 4 (1) | 1 (1) | 5 (1) | 2 (1) | 4 (1) | 1 (1) | 5 (2) | 2 (1) | 5 (1) | 2 (1) |
| N49P6 | 5 (1) | 2 (1) | 4 (1) | 1 (1) | 5 (1) | 2 (1) | 5 (1) | 1 (1) | 4 (2) | 2 (1) | 5 (1) | 2 (1) |
| IOMA | 5 (1) | 2 (1) | 4 (1) | 0 (0) | 6 (1) | 2 (1) | 5 (1) | 0 (0) | 5 (2) | 2 (1) | 6 (1) | 2 (1) |
| N49P7 | 5 (1) | 2 (1) | 4 (1) | 1 (1) | 6 (1) | 2 (1) | 5 (1) | 1 (1) | 5 (2) | 2 (1) | 5 (1) | 2 (1) |
| N60P23 | 4 (1) | 2 (1) | 3 (1) | 1 (1) | 5 (1) | 2 (1) | 5 (1) | 1 (1) | 4 (2) | 2 (1) | 4 (1) | 2 (1) |
| CAP257 | 4 (1) | 1 (1) | 5 (1) | 2 (1) | 7 (1) | 1 (1) | 3 (1) | 1 (1) | 5 (2) | 1 (1) | 3 (1) | 1 (1) |
| 3BNC55 | 5 (1) | 2 (1) | 4 (1) | 0 (0) | 5 (1) | 2 (1) | 5 (1) | 1 (1) | 5 (2) | 2 (1) | 6 (1) | 2 (1) |
| DRVIA7 | 5 (1) | 2 (1) | 4 (1) | 1 (1) | 5 (1) | 2 (1) | 4 (1) | 1 (1) | 6 (2) | 2 (1) | 5 (1) | 2 (1) |

| Antibody | A | | B | | C | | G | | 01_AE | | 07_BC | |
|---|---|---|---|---|---|---|---|---|---|---|---|---|
| | >10 | >50 | >10 | >50 | >10 | >50 | >10 | >50 | >10 | >50 | >10 | >50 |
| 1B2530 | 7 (2) | 2 (1) | 6 (2) | 1 (1) | 5 (1) | 2 (1) | 5 (1) | 1 (1) | 9 (2) | 2 (1) | 6 (2) | 2 (1) |
| VRC06b | 5 (1) | 2 (1) | 4 (1) | 1 (1) | 5 (1) | 2 (1) | 5 (1) | 1 (1) | 5 (2) | 2 (1) | 4 (1) | 2 (1) |
| VRC07 | 4 (1) | 2 (1) | 4 (1) | 1 (1) | 5 (1) | 2 (1) | 4 (1) | 1 (1) | 4 (2) | 2 (1) | 4 (1) | 2 (1) |
| 3BNC117 | 5 (1) | 2 (1) | 4 (1) | 1 (1) | 5 (1) | 2 (1) | 5 (1) | 1 (1) | 6 (2) | 2 (1) | 5 (1) | 2 (1) |
| N6 | 6 (1) | 2 (1) | 4 (1) | 1 (1) | 5 (1) | 2 (1) | 5 (1) | 1 (1) | 7 (2) | 2 (1) | 5 (1) | 2 (1) |
| VRC01 | 4 (1) | 2 (1) | 3 (1) | 1 (1) | 5 (1) | 2 (1) | 4 (1) | 1 (1) | 6 (2) | 2 (1) | 3 (1) | 2 (1) |
| CD4 | 4 (1) | 0 (0) | 3 (1) | 0 (0) | 4 (1) | 1 (1) | 4 (1) | 0 (0) | 13 (2) | 0 (0) | 5 (1) | 0 (0) |
| PGT145 | 9 (3) | 1 (1) | 5 (3) | 1 (1) | 9 (3) | 2 (2) | 7 (3) | 3 (2) | 8 (3) | 5 (3) | 5 (3) | 1 (1) |
| PG16 | 10 (3) | 6 (2) | 5 (2) | 5 (2) | 8 (3) | 5 (2) | 11 (3) | 4 (2) | 11 (3) | 6 (2) | 7 (2) | 5 (2) |
| J033 | 9 (2) | 3 (2) | 5 (2) | 4 (2) | 7 (2) | 6 (2) | 12 (2) | 3 (2) | 9 (2) | 5 (2) | 7 (2) | 3 (1) |
| J038 | 9 (2) | 6 (2) | 5 (2) | 4 (2) | 7 (2) | 6 (2) | 12 (2) | 5 (2) | 9 (2) | 5 (2) | 7 (2) | 5 (2) |
| VRC26.25 | 8 (3) | 2 (1) | 5 (2) | 1 (1) | 8 (3) | 1 (1) | 8 (2) | 1 (1) | 9 (3) | 3 (2) | 6 (2) | 1 (1) |
| PG9 | 8 (2) | 6 (2) | 6 (2) | 4 (2) | 7 (2) | 4 (2) | 11 (2) | 4 (2) | 10 (3) | 6 (2) | 10 (2) | 4 (2) |
| VRC38.01 | 10 (2) | 3 (2) | 6 (2) | 4 (2) | 9 (2) | 3 (2) | 12 (2) | 4 (2) | 9 (2) | 4 (2) | 10 (2) | 3 (1) |
| DH270.5 | 4 (1) | 4 (1) | 6 (1) | 4 (1) | 5 (1) | 5 (1) | 9 (1) | 6 (1) | 7 (1) | 5 (1) | 9 (1) | 7 (1) |
| DH270.6 | 4 (1) | 4 (1) | 6 (1) | 4 (1) | 5 (1) | 5 (1) | 9 (1) | 6 (1) | 7 (1) | 5 (1) | 9 (1) | 7 (1) |
| 10-1074 | 8 (1) | 1 (1) | 9 (1) | 2 (1) | 8 (1) | 2 (1) | 10 (1) | 3 (1) | 9 (1) | 3 (1) | 13 (1) | 3 (1) |
| PGT121 | 5 (1) | 3 (1) | 7 (1) | 3 (1) | 6 (1) | 4 (1) | 8 (1) | 7 (1) | 7 (1) | 4 (1) | 9 (1) | 5 (1) |
| PGT122 | 6 (1) | 3 (1) | 7 (1) | 3 (1) | 6 (1) | 4 (1) | 8 (1) | 6 (1) | 7 (1) | 3 (1) | 10 (1) | 6 (1) |
| PGT128 | 4 (1) | 3 (1) | 6 (1) | 4 (1) | 5 (1) | 5 (1) | 9 (1) | 6 (1) | 7 (1) | 5 (1) | 9 (1) | 6 (1) |
| 438-B11 | 4 (1) | 4 (1) | 6 (1) | 4 (1) | 5 (1) | 4 (1) | 8 (1) | 5 (1) | 7 (1) | 5 (1) | 9 (1) | 7 (1) |
| BG18 | 6 (1) | 2 (1) | 8 (1) | 2 (1) | 8 (1) | 3 (1) | 8 (1) | 2 (1) | 9 (1) | 3 (1) | 10 (1) | 4 (1) |
| PGT135 | 8 (1) | 2 (1) | 10 (1) | 3 (1) | 10 (1) | 3 (1) | 9 (1) | 1 (1) | 9 (1) | 4 (1) | 9 (1) | 4 (1) |
| 8ANC195 | 7 (1) | 2 (1) | 4 (1) | 3 (1) | 5 (1) | 3 (1) | 5 (1) | 3 (1) | 6 (1) | 3 (1) | 5 (1) | 3 (1) |
| 35O22 | 5 (1) | 3 (1) | 4 (1) | 3 (1) | 5 (1) | 4 (1) | 5 (1) | 3 (1) | 4 (1) | 3 (1) | 5 (1) | 3 (1) |
| PGT151 | 6 (2) | 3 (1) | 6 (2) | 4 (1) | 5 (2) | 1 (1) | 5 (2) | 2 (1) | 6 (2) | 2 (1) | 6 (2) | 2 (1) |
| 2G12 | 4 (1) | 0 (0) | 3 (1) | 0 (0) | 3 (1) | 0 (0) | 3 (1) | 0 (0) | 5 (1) | 1 (1) | 5 (1) | 1 (1) |

Table S5: Effective number of overlaping glycan atoms, $\Delta\overline{\mathrm{N}}_{\mathrm{overlap}}$, for CD4bs bNAbs. For each bNAb, the minimum and maximum values are colored blue and red, respectively. Clade B has the most favorable interaction with glycans for most of the bNAbs in this class, indicating the binding of CD4bs will be least hindered by glycans.

| Antibody | Epitope | A | B | C | G | 01_AE | 07_BC |
|---|---|---|---|---|---|---|---|
| VRC01 | CD4bs | 266.50 | 132.60 | 239.92 | 150.91 | 277.30 | 246.76 |
| 3BNC117 | CD4bs | 280.69 | 151.77 | 259.63 | 180.79 | 257.48 | 249.66 |
| VRC07 | CD4bs | 274.55 | 132.29 | 237.26 | 162.27 | 254.57 | 247.09 |
| VRC13 | CD4bs | 643.03 | 422.07 | 682.22 | 558.36 | 648.15 | 722.76 |
| b12 | CD4bs | 209.90 | 404.32 | 329.42 | 330.98 | 543.00 | 619.08 |
| CH103 | CD4bs | 507.42 | 314.50 | 527.20 | 448.71 | 424.55 | 510.36 |
| HJ16 | CD4bs | 225.95 | 312.41 | 336.90 | 231.63 | 208.38 | 271.58 |
| NC-Cow1 | CD4bs | 149.64 | 69.69 | 134.36 | 73.31 | 162.37 | 117.18 |
| VRC02 | CD4bs | 392.86 | 238.48 | 397.76 | 302.89 | 544.26 | 409.80 |
| 8ANC131 | CD4bs | 402.01 | 245.30 | 414.29 | 308.15 | 586.56 | 410.67 |
| VRC-PG20 | CD4bs | 262.67 | 137.22 | 244.29 | 166.00 | 249.12 | 241.34 |
| VRC-CH31 | CD4bs | 343.29 | 187.27 | 308.13 | 226.18 | 315.58 | 307.01 |
| 12A21 | CD4bs | 321.31 | 184.46 | 299.17 | 223.64 | 374.14 | 284.50 |
| NIH45-46 | CD4bs | 253.61 | 125.12 | 231.97 | 146.67 | 247.31 | 235.97 |
| VRC-PG04 | CD4bs | 97.62 | 4.09 | 80.49 | 23.34 | 111.29 | 74.30 |
| CH235.9 | CD4bs | 278.69 | 139.37 | 257.50 | 169.17 | 342.67 | 263.31 |
| CH235.12 | CD4bs | 272.32 | 132.62 | 239.88 | 162.16 | 275.12 | 241.49 |
| 179NC75 | CD4bs | 187.41 | 231.06 | 270.43 | 216.62 | 223.35 | 237.21 |
| 2411a | CD4bs | 293.13 | 151.09 | 254.09 | 175.37 | 279.19 | 265.96 |
| N49P6 | CD4bs | 294.22 | 155.16 | 264.37 | 187.17 | 286.85 | 265.09 |
| N49P7 | CD4bs | 296.95 | 162.29 | 272.58 | 197.45 | 284.73 | 267.05 |
| N60P23 | CD4bs | 265.17 | 135.95 | 241.32 | 166.41 | 247.70 | 235.36 |
| CAP257 | CD4bs | 185.40 | 160.71 | 207.46 | 154.89 | 233.94 | 215.41 |
| 3BNC55 | CD4bs | 137.22 | 42.47 | 122.96 | 54.03 | 135.50 | 137.12 |
| DRVIA7 | CD4bs | 286.16 | 146.86 | 257.17 | 169.07 | 268.95 | 263.93 |
| 1B2530 | CD4bs | 139.38 | 54.17 | 129.98 | 62.60 | 203.01 | 135.89 |
| VRC06b | CD4bs | 280.51 | 140.71 | 243.28 | 171.12 | 261.61 | 250.63 |
| N6 | CD4bs | 318.99 | 168.99 | 292.64 | 208.20 | 323.40 | 282.00 |
| BG24 | CD4bs | 287.40 | 152.70 | 254.86 | 176.82 | 264.23 | 269.96 |
| IOMA | CD4bs | 255.56 | 139.04 | 252.03 | 163.32 | 248.88 | 264.91 |

Table S6: Effective number of overlaping glycan atoms, $\Delta\overline{\mathrm{N}}_{\mathrm{overlap}}$, for V1/V2 apex (V2), V3-glycan (V3), gp120/gp41 interface (Interface), Fusion Peptide (FP) bNAbs. For each bNAb, the minimum and maximum values are colored blue and red, respectively. Clade B has the most favourable interactions for all of V1/V2 apex bNAbs. Depending on the bNAb, either clade 01_AE or 07_BC has the least favorable glycan interaction for V3-glycan class bNAbs. 2G12, which only target, has many overlapping glycan atoms in clades 01_AE or 07_BC that are not known to be important for its binding or recognition.

| Antibody | Epitope | A | B | C | G | 01_AE | 07_BC |
|---|---|---|---|---|---|---|---|
| PG9 | V1V2 apex | 177.34 | -47.16 | 105.18 | 112.05 | 228.31 | 68.66 |
| PG16 | V1V2 apex | 220.73 | 77.61 | 123.50 | 203.91 | 316.99 | 103.60 |
| PGT145 | V1V2 apex | 12.28 | -78.89 | -53.30 | 16.11 | 144.87 | -49.38 |
| J033 | V1V2 apex | 152.47 | 82.27 | 179.62 | 239.13 | 329.67 | 123.31 |
| J038 | V1V2 apex | 174.53 | 90.34 | 194.33 | 264.39 | 364.58 | 142.12 |
| VRC26.25 | V1V2 apex | 213.45 | 59.53 | 78.21 | 148.22 | 259.31 | 149.71 |
| VRC38.01 | V1V2 apex | 68.65 | -45.52 | 88.57 | 144.64 | 147.12 | 0.29 |
| DH270.5 | V3-glycan | 69.36 | 77.52 | 107.37 | 207.97 | 422.57 | 334.64 |
| DH270.6 | V3-glycan | 77.95 | 80.76 | 112.62 | 209.74 | 409.13 | 347.78 |
| 10-1074 | V3-glycan | 126.29 | 185.71 | 178.66 | 233.56 | 320.91 | 409.19 |
| PGT121 | V3-glycan | 231.18 | 388.03 | 433.85 | 481.25 | 406.92 | 653.46 |
| PGT122 | V3-glycan | 48.89 | 109.29 | -9.35 | 120.88 | 308.34 | 338.60 |
| PGT128 | V3-glycan | 35.30 | 54.58 | -155.00 | -38.73 | 338.68 | 85.91 |
| 438-B11 | V3-glycan | 257.83 | 473.76 | 486.02 | 558.33 | 487.01 | 672.40 |
| BG18 | V3-glycan | 211.22 | 190.51 | 233.55 | 250.47 | 437.20 | 425.43 |
| PGT135 | V3-glycan | 192.79 | 159.35 | 184.30 | 4.75 | 420.07 | 233.66 |
| 8ANC195 | Interface | -124.30 | -138.70 | -108.74 | -110.45 | -66.65 | -106.90 |
| 35O22 | Interface | 269.83 | 57.97 | 121.69 | 99.88 | 60.60 | 35.32 |
| 2G12 | Glycan | 20.38 | 18.51 | 16.94 | 21.41 | 172.75 | 125.71 |
| PGT151 | Fusion Peptide | 279.60 | 391.23 | 181.62 | 289.33 | 302.66 | 261.60 |

### Interaction of PGT151 with Clade B Glycans

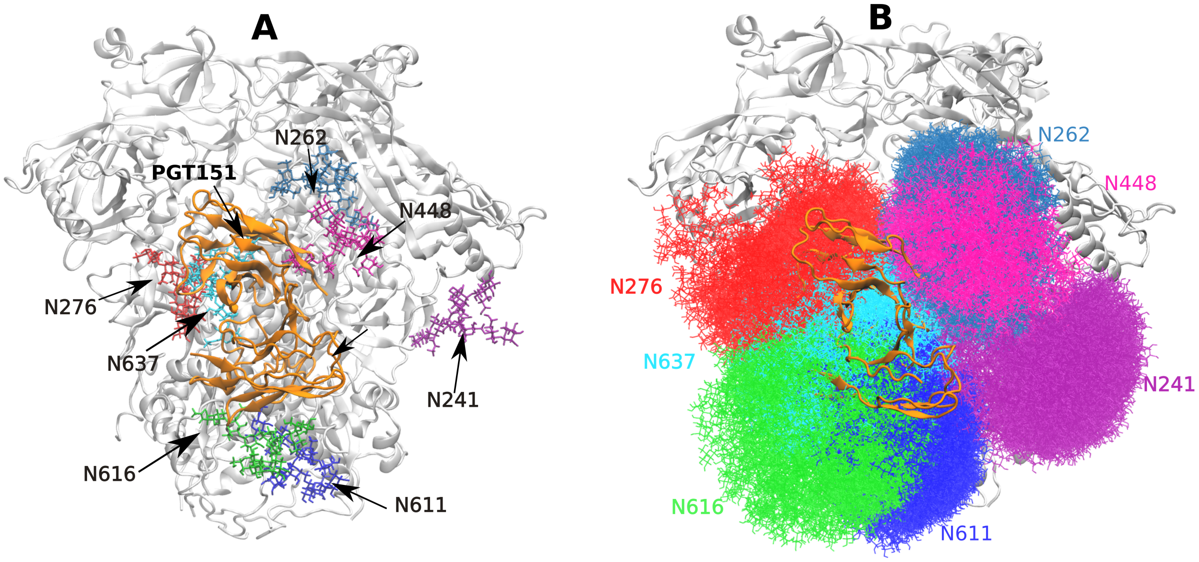


Figure S5: (A) Structure of Clade B trimer (white) bound to PGT151 (orange) bNAb. Modeled Man9 glycans at key PNGS are also shown. (B) 300 glycan conformers at each PNGS generated using GlycoSHIELD are shown to illustrate PGT151-glycan overlap. PGT151 has significant overlap with glycans at N276, N616, and N637 of one protomer and aslo has contacts with glycans at N262 and N448 of the adjacent protomer. Only a very few atoms of N241 glycans are near the bound PGT151 bNAb.

Table S7: Values of total epitope shielding, $\mathcal{S}$, for various bNAbs. $\mathcal{S}$ gives a measure of how the protein epitopes of bNAbs are shielded by glycans.

| Antibody | Epitope | A | B | C | G | 01_AE | 07_BC |
|---|---|---|---|---|---|---|---|
| VRC13 | CD4BS | 54.21 | 47.55 | 55.32 | 51.58 | 39.91 | 54.40 |
| b12 | CD4BS | 30.09 | 29.48 | 34.19 | 26.90 | 25.97 | 38.69 |
| CH103 | CD4BS | 45.75 | 51.97 | 52.61 | 40.79 | 31.55 | 41.81 |
| BG24 | CD4BS | 39.86 | 28.75 | 38.76 | 32.45 | 20.32 | 28.20 |
| HJ16 | CD4BS | 13.29 | 12.66 | 26.53 | 14.02 | 8.23 | 12.21 |
| NC-Cow1 | CD4BS | 36.14 | 32.97 | 40.04 | 38.84 | 24.65 | 38.28 |
| 8ANC131 | CD4BS | 25.28 | 22.35 | 20.22 | 23.96 | 19.48 | 29.58 |
| 12A21 | CD4BS | 22.28 | 30.84 | 35.17 | 32.99 | 18.51 | 16.67 |
| NIH45-46 | CD4BS | 69.89 | 71.96 | 77.05 | 67.46 | 52.59 | 74.21 |
| VRC-PG04 | CD4BS | 64.47 | 52.88 | 56.44 | 58.15 | 47.05 | 58.49 |
| 2411a | CD4BS | 58.84 | 55.78 | 64.39 | 59.91 | 49.33 | 63.03 |
| N49P6 | CD4BS | 63.38 | 52.32 | 61.29 | 55.18 | 47.25 | 60.68 |
| IOMA | CD4BS | 21.86 | 20.07 | 25.69 | 25.37 | 11.27 | 17.17 |
| N49P7 | CD4BS | 57.94 | 44.52 | 51.06 | 54.06 | 43.76 | 53.67 |
| N60P23 | CD4BS | 56.14 | 44.80 | 52.75 | 45.40 | 39.94 | 54.17 |
| DRVIA7 | CD4BS | 54.06 | 54.48 | 51.80 | 45.72 | 40.15 | 50.89 |
| 1B2530 | CD4BS | 27.69 | 21.85 | 22.03 | 29.67 | 20.30 | 30.93 |
| 3BNC117 | CD4BS | 82.81 | 74.65 | 89.21 | 82.06 | 61.11 | 85.23 |
| N6 | CD4BS | 75.08 | 73.76 | 64.73 | 68.52 | 50.93 | 70.62 |
| VRC01 | V1V2 apex | 70.87 | 74.63 | 86.27 | 73.65 | 55.70 | 72.51 |
| PGT145 | V1V2 apex | 14.63 | 25.09 | 20.42 | 16.11 | 42.02 | 32.56 |
| PG16 | V1V2 apex | 15.30 | 26.80 | 23.14 | 17.66 | 42.02 | 32.56 |
| J038 | V1V2 apex | 42.10 | 47.44 | 38.32 | 44.23 | 64.93 | 59.96 |
| VRC26.25 | V1V2 apex | 1.93 | 11.00 | 11.00 | 7.31 | 21.79 | 19.69 |
| PG9 | V1V2 apex | 15.30 | 26.80 | 23.14 | 17.66 | 42.02 | 32.56 |
| VRC38.01 | V3-glycan | 11.10 | 19.46 | 22.51 | 12.22 | 43.79 | 23.59 |
| 10-1074 | V3-glycan | 40.18 | 49.70 | 68.14 | 74.37 | 44.53 | 58.56 |
| PGT121 | V3-glycan | 49.67 | 59.83 | 76.64 | 81.18 | 34.26 | 58.56 |
| PGT122 | V3-glycan | 10.16 | 11.90 | 13.66 | 14.49 | 12.15 | 3.93 |
| PGT128 | V3-glycan | 46.44 | 58.46 | 73.98 | 74.19 | 61.32 | 80.94 |
| 438-B11 | V3-glycan | 20.97 | 15.35 | 18.59 | 13.67 | 13.50 | 20.06 |
| BG18 | V3-glycan | 51.35 | 58.38 | 69.38 | 51.39 | 29.07 | 41.12 |
| PGT135 | V3-glycan | 25.70 | 33.11 | 28.49 | 7.51 | 24.31 | 25.39 |
| 8ANC195 | Interface | 61.04 | 54.66 | 43.86 | 60.50 | 42.60 | 45.31 |
| 35O22 | Interface | 16.93 | 13.48 | 24.44 | 13.55 | 18.56 | 16.14 |
| PGT151 | Fusion Peptide | 32.97 | 46.35 | 31.03 | 35.14 | 36.85 | 31.66 |
| 2G12 | glycan | 36.24 | 21.53 | 17.03 | 21.00 | 28.34 | 26.85 |

### Native-like vs Man9 glycosylation

#### Total Accessible Surface Area

Table S8 shows how total ASA with Man9 glycosylation compares against the native-like glycosylation. Although there is a variation in the absolute value of ASA for a given probe, the change in ASA with probe radii is comparable for both Man9 and Native-like glycosylation.

Table S8: Total accessible surface area (Å$^2$) for with Man9 and native-like fully glycosylated HIV-1 Envs for clade A with a probe radii of 1.4, 7.2, and 10.0Å. see Table 1 for site-specific glycoforms used in native-like glycosylation.

| | $SASA_{tot}$ with Man9 glycosylation | | | $SASA_{tot}$ with native-like glycosylation | | |
|---|---|---|---|---|---|---|
| probe size (Å) | 1.4 | 7.2 | 10.0 | 1.4 | 7.2 | 10.0 |
| | 66296.3 | 22900.5 | 20100.7 | 65054.3 | 21021.6 | 18214.5 |

### AA Sequences of Clades

Following sections shows the AA sequences of each clade aligned to the Hxb2 sequence using clustalo software[2]. All the modeled missing residues (using Modeller software) are colored either blue or green, where green color is used for the PNGS. All other PNGS are colored red.

#### 5FYL (Clade A): 31-664

**gp120:**

--AENLWVTVYYGVPVWKDAETTLFCASDAKAYETEKHNVWATHACVPTDPN PQEIHLEN

VTEEFNMWKNNMVEQMHTDIISLWDQSLKPCVKLTPLCVTLQCTNVTNNITDD--------MRG

ELKNCSFNMTTELRDKKQKVYSLFYRLDVVQINENQGNRSNNSNKEYRLINCNTSAITQACP

KVSFEPIPIHYCAPAGFAILKCKDKKFNGTGPCPSVSTVQCTHGIKPVVSTQLLLNGSLAEEE

VMIRSENITNNAKNILVQFNTPVQINCTRPNNNTRKSIRI--GPGQAFYATGDIIGDIRQAHCNV

SKATWNETLGKVVKQLRKHFGNNTIIRFANSSGGDLEVTTHSFNCGGEFFYCNTSGLFNSTW

ISNTSVQGSNSTGS-NDSITLPCRIKQIINMWQRIGQAMYAPPIQGVIRCVSNITGLILTRDGGST

NSTTETFRPGGGDMRDNWRSELYKYKVVKIEPLGVAPTRCKRRVVGRRRRR

**gp41:**

AVGIGAVFLGFLGAAGSTMGAASMTLTVQARNLLSGIVQQQSNLLRAPEAQQHLLKLTVWG
IKQLQARVLAVERYLRDQQLLGIWGCSGKLICCTNVPWNSSWSNRNLSEIWDNMTWLQWDK
EISNYTQIIYGLLEESQNQQEKNEQDLLALD

#### 5FYK (Clade B) : 31-672

**gp120:**

--VEKLWVTVYYGVPVWKEATTTLFCASDAKAYDTEVHNVWATHACVPTDPN PQEVVLENV
TEHFNMWKNNMVEQMQEDIISLWDQSLKPCVKLTPLCVTLNCKDVNATNTTNDSEGT--MER
GEIKNCSFNITTSIRDKVQKEYALFYKLDVVPIDNNNTSYRLISCDTSVITQACPKISFEPIPIHYC
APAGFAILKCNDKTFNGKGPCKNVSTVQCTHGIRPVVSTQLLLNGSLAEEEVVIRSDNFTNNAK
TIIVQLKESVEINCTRPNNNTRKSIHI--GPGRAFYTTGEIIGDIRQAHCNISRAKWNDTLKQIVIK
LREQFEN-KTIVFNHSSGGDPEIVMHSFNCGGEFFYCNSTQLFNSTWNNNTEGSNNTE----GNTI
TLPCRIKQIINMWQEIGKAMYAPPIRGQIRCSSNITGLLLTRDGCINENGTEIFRPGGGDMRDNW
RSELYKYKVVKIEPLGVAPTKCKRRVVGRRRR

**gp41:**

AVGIGAVFLGFLGAAGSTMGAASMTLTVQARLLLSGIVQQQNNLLRAPEAQERMLQLTVWGIK
QLQAR VLAVERYLGDQQLLGIWGCSGKLICCTAVPWNASWSNKSLDRIWNNMTWMEWEREI
DNYTSEIYTLIEESQNQQEKNEQELLELDGGLEVLFQ

#### 8SAU (Clade C): 31-664

**gp120:**

--AENLWVTVYYGVPVWKEAKTTLFCASDARAYEKEVHNVWATHACVPTDPSPQELVLGNVTE

NFNMWKNDMVDQMHEDIISLWDQSLKPCVKLTPLCVTLICSNATVKNG----------TVEEMKNCSF
NTTTEIRDKEKKEYALFYKPDIVPLSETNNTSEYRLINCNTSACTQACPKVTFEPIPIHYCAPAGY
AILKCNDETFNGTGPCSNVSTVQCTHGIRPVVSTQLLLNGSLAEKEIVIRSENLTNNAKIIIVHLHT
PVEIVCTRPNNNTRKSVRI--GPGQTFYATGDIIGDIKQAHCNISEEKWNDTLQKVGIELQKHFP-
NKTIKYNQSAGGDMEITTHSFNCGGEFFYCNTSNLFNGTYNGTYISTNSSA--NSTSTITLQCRIKQ
IINMWQGVGRCMYAPPIAGNITCRSNITGLLLTRDGGTNSNETETFRPAGGDMRDNWRSELYKY
KVVKIEPLGVAPTRCKRRV------

**gp41:**

AVGIGAVFLGFLGAAGSTMGAASMTLTVQARNLLSG--------------------TVWGIKQLQARVLAVERY
LRDQQLLGIWGCSGKLICCTNVPWNSSWSNRNLSEIWDNMTWLQWDKEISNYTQIIYGLLEESQ
NQQEKNEQDLLALD

#### 8GPI (CRF01_AE): 33-664

**gp120:**

----NLWVTVYYGVPVWRDADTTLFCASDAKAHVPEAHNVWATHACVPTDPNPQEIPLENVTE
NFNMWKNNMVEQMQEDVISLWDQSLKPCVKLTPLCVTLNCTKANLTHNTTNDKNGTGNITD
EVK--------IGNITDEVKNCTFNMTTEIRDKQQKVHALFYALDIVQMKENGSEYRLISCNTSVIKQ
ACPKISFDPIPIHYCAPAGYAILKCNDKKFNGTGPCKNVSTVQCTHGIKPVVSTQLLLNGSLAE
EEIIIRSENLTNNAKNIIVHLNKSVSISCTRPSNNTRTSIRI--GPGQMFYRTGDIIGDIRKAYCELNG
TEWNETLNKVTEKLKEHF--NKTIVFQPPSGGDLETTMHHFNCRGEFFYCNTTKLFNT-------
--KNGTREEFNGTIILPCRIKQIVNMWQGVGQAMYAPPISGIINCTSNITGIILTRDGGNGNTTDE
TFRPGGGNIKDNWRSELYKYKVVQIEPLGIAPTRCKRR--------

**gp41:**

--------GFLGAAGSTMGAASITLTVQARQL-------------LSGNPDWLPDMTVWGIKQLQARVLAVER
YLKDQKFLGLWGCSGKIICCTNVPWNSTWSNKSYEEIWNNMTWIEWEKEISNYTNRIYDLLTE
SQNQQERNEKDLLELD

#### 8GPJ (CRF07_BC): 33-660

**gp120:**

----NLWVTVYYGVPVWKEATTTLFCASDAKAYDTEVHNVWATHACVPTDPNPQEMVLENV

TENFNMWKNEMVNQMHEDVISLWDQSLKPCVKLTPLCVTLDCTTVNSNSSSNSSNSSGNSN

STLEDM-----------------QEMKNCSFNTTTELRDKKQKVYALFYKLDIVPLSNNSSEYRLINCNTS

AITQACPKVSFDPIPIHYCTPAGYALLKCNDKRFNGTGPCHNVSTVQCTHGIKPVVSTQLLL

NGSLAEKEIIVRSENLTNNVKTIIVHLNKSVEIVCTRPGNNTRKSIRI-- GPGQTFYATGDIIGD

IRQAHCNISRGDWEETLHNVRKNLAEHFQ-NKTIQFASSSGGDLEITTHSFNCRGEFFYCNTS

GLFNST----YMPNSTFNGTESNLTITIPCRIKQIINMWQEVGRAMYAPPIAGNITCKSNITGLLL

VRDGGKESNSTEIFRPGGGDMRDNWRSELYKYKVVEIKPLGVAPTECKRR--------

**gp41:**

-------LGFLGVAGSTMGAASVALTVQARQLLSG------NPDW-------LPDMTVWGIKQLQTRVLAI

ERYLKDQQLLGIWGCSGKLICCTAVPWNSSWSNKSQTEIWNNMTWMQWDEEISNYTATIYR

LLEVSQNQQERNEKDL

LALD

#### 5FYJ (Clade G): 29-672

**gp120:**

ALAGDLWVTVYYGVPVWEDADTTLFCASDAKAYSTESHNVWATHACVPTDPNPQEIPLKN

VTENFNMWKNNMVEQMHEDIISLWDESLKPCVKLTPLCVTLICTNVTSNSTNSTNGVTNNS

T---VDYREQLKNCSFNITTEIRDKQRKEYALFYRLDIVPINDNEKNDTYRLINCNVSTIKQACP

KVTFDPIPIHYCAPAGFAILKCRDKKFNGTGPCKNVSTVQCTHGIKPVISTQLLLNGSLAEGD

IMIRSENITDNAKTIIVQLKTAVNITCTRPSNNTRKSIRF--GPGQAFYATDEIIGDIRQAHCNISK

TEWEDMKRNVSDKLKALFNN-KTIIFKSSSGGDLEITTHSFNCRGEFFYCNTSGLFNTSGLFN-

-----NNSNDSSGNITLPCKIKQIVRMWQRVGQAMYAPPIAGNITCRSRITGLLLVRDGCKSNETN GTETFRPAGGDMRDNWRSELYKYKVVKIKPLGVAPTRCRRRVVGRRRRR

**gp41:**

AVGMGAVLLGFLGAAGSTMGAASITLTVQVRQLLSGIVQQQSNLLRAPEAQQHLLQLTVWGI KQLQARVLAVERYLQDQQLLGIWGCSGRLICCTNVPWNASWSNKSYSEIWDNLTWVEWERE ISNYTQHIYNLLQESQNQQEKNEQDLLALDKGGGLVPR

### References


(1) Chakraborty, S.; Berndsen, Z. T.; Hengartner, N. W.; Korber, B. T.; Ward, A. B.; Gnanakaran, S. Quantification of the Resilience and Vulnerability of HIV-1 Native Glycan Shield at Atomistic Detail. *iScience* **2020**, *23*, 101836.

(2) Sievers, F.; Higgins, D. G. Clustal omega. *Current protocols in bioinformatics* **2014**, *48*, 3–13.
